# Nucleoporin Nsp1 surveils the phase state of FG-Nups

**DOI:** 10.1101/2023.03.31.535084

**Authors:** Tegan A. Otto, Tessa Bergsma, Maurice Dekker, Sara N. Mouton, Paola Gallardo, Justina C. Wolters, Anton Steen, Patrick R. Onck, Liesbeth M. Veenhoff

## Abstract

Transport through the NPC relies on intrinsically disordered FG-Nups forming a selective barrier. Away from the NPC, FG-Nups readily form condensates and aggregates, and we address how this behavior is surveilled in cells. FG-Nups, including Nsp1, together with nuclear transport receptor Kap95, form a native cytosolic condensate in yeast. In aged cells this condensate disappears as cytosolic Nsp1 levels decline. Biochemical assays and modeling show that Nsp1 is a modulator of FG-Nup liquid-liquid phase separation, promoting a liquid-like state. Nsp1’s presence in the cytosol and condensates is critical, as a reduction of cytosolic levels in young cells induces NPC assembly and transport defects and a general decline in protein quality control, all quantitatively mimicking aging phenotypes. Excitingly, these phenotypes can be rescued by cytosolic Nsp1. We conclude that Nsp1 is a phase state regulator that surveils FG-Nups and impacts general protein homeostasis.

**Highlights:**

- Nups form native cytosolic condensates
- Nsp1 reduction mimics NPC aging phenotypes
- Nsp1 acts as phase state modulator of FG-Nups
- Nsp1 shares surveillance function with classical chaperones

## Introduction

Nuclear pore complexes (NPCs) represent a major hub in the eukaryotic cell by forming a selective barrier for the import and export of RNAs and proteins between the cytosol and nucleus^1, 2^. It is however not trivial to assemble and maintain this essential protein complex. The challenges are related to its impressive size – NPCs are composed of more than 500 proteins^3–7^ making assembly an error-prone process^8, 9^. Also, the existence of long-lived subunits within the NPC makes NPCs susceptible to the accumulation of damage over time^10–12^. Finally, large parts of the NPC are built from intrinsically disordered proteins (IDPs)^13^, a class of proteins that readily self-associate and potentially aggregate^14–20^. The assembly and maintenance of a protein complex as large and complex as the NPC is thus tremendously challenging for a cell.

In mitotically aging yeast cells NPC components are no longer present in the correct amounts (loss of stoichiometry)^21–23^, components involved in NPC quality control decline in abundance and misassembled NPCs accumulate^22^. Also, nuclear transport becomes dysfunctional in aging cells and in aggregation pathologies^11, 24–31^. For cells challenged with aggregation prone peptides, such as in aggregation pathologies, a main challenge may be to guard the disordered proteins of the NPC. This follows from evidence that the nuclear transport machinery modulates the phase state and toxicity of IDPs related to neurodegenerative diseases^25, 27–29, 31, 32^. These newly found vulnerabilities of eukaryotic cells inspire to study the mechanisms that control the quality of NPCs, and especially the ID NPC components.

The ID NPC components (nucleoporins, nups) are characterized by repeats of phenylalanine and glycine (FG) and hence categorized as FG-Nups^13^.They serve to form the selective permeability barrier in NPCs together with nuclear transport receptors (NTRs)^2, 13, 33–36^. FG-Nups and NTRs are also critical in the assembly of NPCs^37, 38^. In isolation, the FG-Nups, like other IDPs, can phase separate into condensates and form amyloids ^39–50^. Within the NPC, the FG-Nups are however anchored and surrounded by a mix of FG-repeat sequences that may be evolutionary finetuned to assure dynamicity and counter phase separation and aggregation. Also, the large numbers of NPC resident NTRs likely assist in preventing aggregation of FG-Nups^40^. While such mechanisms may suffice in mature NPCs in healthy young cells, additional mechanisms may be needed to guard the FG-Nups in aging or under conditions of stress, such as exposure to aggregation prone proteins. Surveillance is certainly needed in the timeframe between synthesis and incorporation in the NPC and the recently discovered mechanisms include local translation at NPCs, co-translation of subcomplexes^51, 52^, NTR binding^38, 53^ and surveillance by molecular chaperones^54, 55^. Additionally, nup condensates have been described to serve as assembly platforms^56, 57^.

To address the question how FG-Nups may be susceptible to aggregation in aging, we focus on Nsp1. Nsp1, one of the first nups identified^58^, is an abundant central channel FG-Nup present in 48 copies per NPC^59^. It forms two heterotrimeric subcomplexes, the Nsp1-Nup49-Nup57 complex in the equator of the NPC central channel, and the Nsp1-Nup82-Nup159 complex facing the cytosol^59–62^ (Figure 1A, Figure S1A). We reason Nsp1 is a good target to address how FG-Nups may be at risk in aging, because its abundance decreases significantly in aged yeast cells^21, 23^ (Figure 1B, Figure S1B). Also, a cytosolic pool of Nsp1, forming an ER-associated daughter specific focus during cell division, has already been implicated in pore inheritance and pore assembly in the daughter cell^63, 64^. Compared to other FG-Nups, Nsp1 has a unique amino acid sequence with almost perfectly repeated FG-repeat sequences interspaced with charged spacers (Figure S1C). Such a sequence is exemplary of an organization in “stickers and spacers” in which the spacers tune the solubility and regulate the phase behavior of IDPs^14, 15, 50^. Indeed, the first experiments showing FG-Nup hydrogels were actually done with the FG-domain of Nsp1^39^.

**Figure 1.**
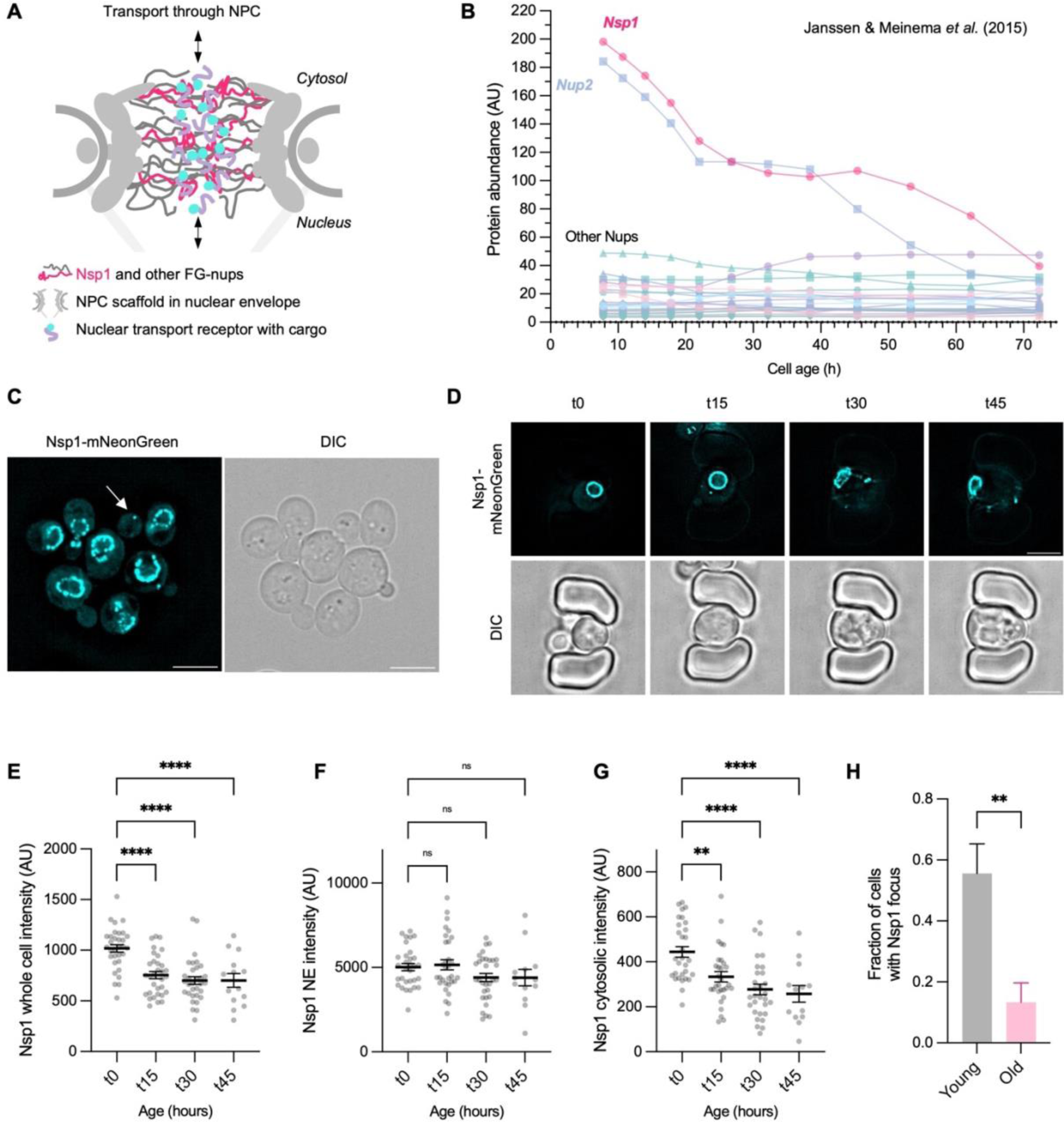
Nsp1 localizes in three distinct pools and decreases with aging. (A) Cartoon of the Nuclear Pore Complex. (B) Nup protein levels in aging. Data from Janssen & Meinema *et al.* (2015)^21^ (C) Representative image of cells endogenously expressing mNeonGreen-tagged Nsp1. White arrow indicates the daughter specific Nsp1 focus. Scale bar, 5µm (D) Images of an aging cell expressing endogenously Nsp1-mNeonGreen as observed in a microfluidic chip over 48h. Scale bar, 5µm (E) Quantification of endogenous Nsp1-mNeongreen whole cell signal in aging cells. Graph shows mean ± SEM of >30 cells (n=2). (F) Quantification of endogenous Nsp1-mNeongreen signal at the NE in aging cells. Graph shows mean ± SEM of >30 cells (n=2). (G) Quantification of cytosolic Nsp1-mNeongreen signal in aging mother cells. Graph shows mean ± SEM of >30 cells (n=2). (H) Frequency of daughter specific Nsp1 focus in young and batch-aged cells (20 hours). Graph shows mean ± SEM of >30 cells per condition. ns: not significant, **p < 0.01, ****p < 0.0001

This study identifies the nucleoporin Nsp1 as an important modulator for FG-Nup phase state control prior to NPC assembly and shows that a cytosolic pool of Nsp1 can rescue age-related phenotypes including and beyond NPC biology. We propose Nsp1 shares a surveillance function with classical chaperones in maintaining protein homeostasis.

## Results

### Nsp1 localizes in three distinct pools, and decreases with aging

To better define the abundance and localization of Nsp1 we endogenously tagged Nsp1 with a fast-folding and bright mNeonGreen fluorescent protein^65^. Nsp1 indeed shows a punctate rim staining at the NE that is typical for nucleoporins, but also a cytosolic pool is observed (Figure 1C, Figure S2AB). In about 65% of emerging daughter cells (S-phase of cell cycle; bud size 2-3.5 micron) a bright focus is visible (Figure 1C, 2F and previously described in ^63^), which we refer to as the daughter specific Nsp1 focus. Following cells for 48 hours (replicative age around 10) in a microfluidic chip^66^ allowed us to observe changes in the levels of the different Nsp1-mNeonGreen pools (Figure 1D, Figure S2E). Consistent with previous data^21, 23^, the whole cell intensity of Nsp1 decreases in aging (Figure 1E). Whereas the NE pool shows no significant changes, we observe a decrease in the disperse cytoplasmic Nsp1 pool (Figure 1FG, Figure S2CD). To assess if the appearance of the daughter specific focus changes in aging we resorted to a 20h batch-aging experiment which allows imaging more cells than in a microfluidic chip. This analysis shows that the daughter specific Nsp1 focus is less frequently observed in emerging daughters of mid-aged cells (Figure 1H). Distinct from the daughter specific Nsp1 foci, we occasionally also observe foci in the cytosol of mother cells. With increased age these latter foci, which probably represent non-functional forms of the protein, become more frequent (Figure S2F). Together, these results show that Nsp1 is present in three different pools: it is localized in the NPC, in a daughter specific focus and dispersed in the cytoplasm. Total Nsp1 levels reduce with age^21, 22^ and here we specifically show a decrease in the cytoplasmic pool and the frequency of the daughter specific focus.

**Figure 2.**
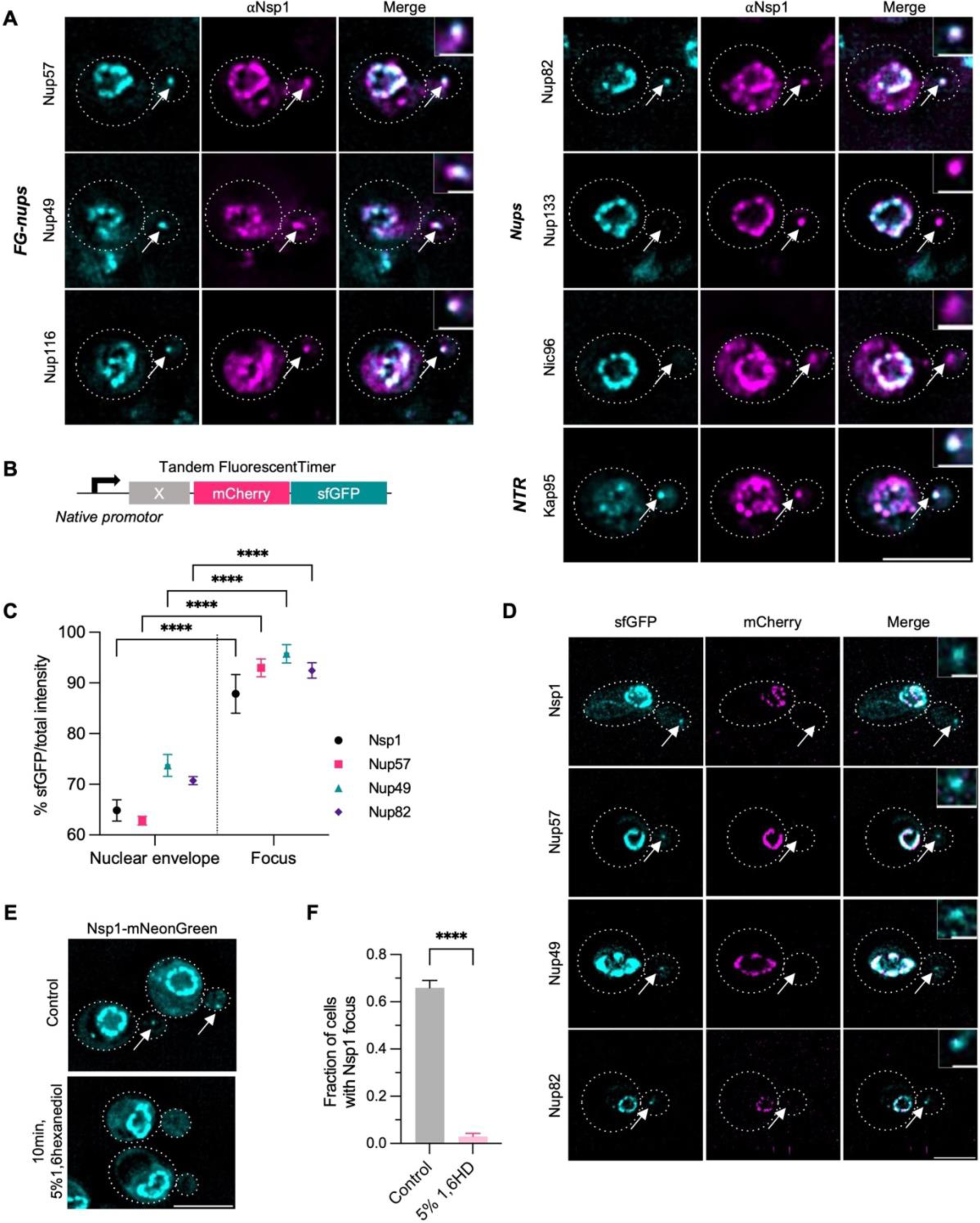
Characterization of native cytosolic Nsp1 condensate. (A) Representative immunofluorescence images of Nup-GFP with Nsp1-antibody. White arrows indicate daughter specific Nsp1 foci. Dashed lines outline the cells. Inset represents a zoom in of the Nsp1 focus. Scale bar, 5µm. Scale bar inset, 1µm. (B) Graphical representation of tandem-fluorescent timer^68^. sfGFP: superfolder GFP (C) Percentage new protein (sfGFP) over total protein (sfGFP+mCherry signal) comparing the ratio at the nuclear envelope and in the daughter specific Nsp1 focus. Graph shows mean ± SEM of >10 cells per condition (n=3). (D) Representative images of cells expressing endogenous tandem fluorescent timer-fusions of Nsp1, Nup57, Nup49 and Nup82. White arrows indicate daughter specific Nsp1 foci. Scale bar, 5µm. Scale bar inset, 1µm. (E) Representative images of Nsp1-mNeongreen cells with and without treatment of 1,6-hexanediol (5%, 10 min). Scale bar, 5µm. (F) Fraction of cells with an Nsp1 focus with and without treatment with 1,6-hexanediol. Graph shows mean ± SEM of 150 cells per condition (n=3). ****p < 0.0001

### A native cytosolic condensate of Kap95 and newly synthesized nups

Previous work using a RITE-cassette approach has already shown that the daughter specific Nsp1 focus contained newly synthesized Nsp1^63^ and Nup57^67^. Here, we assessed colocalization of several other endogenously GFP-tagged nups with the daughter specific Nsp1 focus using immunofluorescence with an Nsp1-antibody. Colocalization of the Nsp1-antibody and the GFP signal was found for Nup57 (57% overlap between both signals), Nup49 (81%), Nup82 (53%) and Nup116 (51%) (Figure 2A, Figure S3A). These nups form subcomplexes in the mature NPC (Figure S1A)^59^ and/or are co-translationally assembled (e.g. Nsp1 and Nup57, Nup82 and Nup116)^51, 52^. Colocalization is rarely observed for the scaffold nups Nup133 (5%) and Nic96 (1%). The NTR Kap95 also showed strong overlap with the Nsp1 signal in the focus (79%) (Figure 2A, Figure S3A). A tandem-fluorescent timer^68^ approach showed that the Nsp1 focus contains primarily newly synthesized Nsp1 as well as new Nup57, Nup49 and Nup82, while the NPC resident pools of these proteins are older (sfGFP/total fluorescence in the focus being 88-96% and at the NE 62-70%) (Figure 2B-D).

We next asked if the daughter specific Nsp1 focus is a liquid-liquid phase separated condensate. The focus is too small to assess using FRAP, and hence we studied if exposure of cells to 1,6-hexanediol would dissolve it. 1,6-Hexanediol is an aliphatic alcohol and 10 min exposure to 5% 1,6-hexanediol provides a surprisingly specific disruption of the interactions between FG-Nups and NTRs^69^ and between FG-Nups ^41, 43–46^, and generally dissolves liquid-like compartments^70^. Indeed, the exposure to 1,6-hexanediol reduced the frequency of observing the focus in the emerging daughter from 65% to less than 3%, strongly indicating that the Nsp1 focus is a phase-separated, liquid-like condensate (Figure 2EF).

Together these results show that the native daughter specific Nsp1 focus is a biological condensate consisting of newly synthesized nups, and the NTR Kap95.

### Nsp1 and Kap95 modulate FG-Nup phase state *in vitro* and in cells

The liquid-like or gel-like states generally represent the functional states of biological condensates, while the evolution into an aggregated state is often associated with a loss of function or even gain of toxicity^71–76^. We studied how the native Nsp1 condensate may be stabilized in a functional state. FG-Nups engage with NTRs^2^, and therefore we tested whether purified Kap95 is able to alter the phase state of *in vitro* formed condensates of the FG domain of Nup116 (Nup116_1-725,_ Nup116FG from hereon). Both Kap95 and Nup116 are present in the daughter specific Nsp1 focus. Visual inspection of the condensates shows that the presence of equimolar amounts of Kap95 alters their appearance from a mixture of differently shaped particles to a mixture of rounder particles of varying size (Figure 3A).

**Figure 3.**
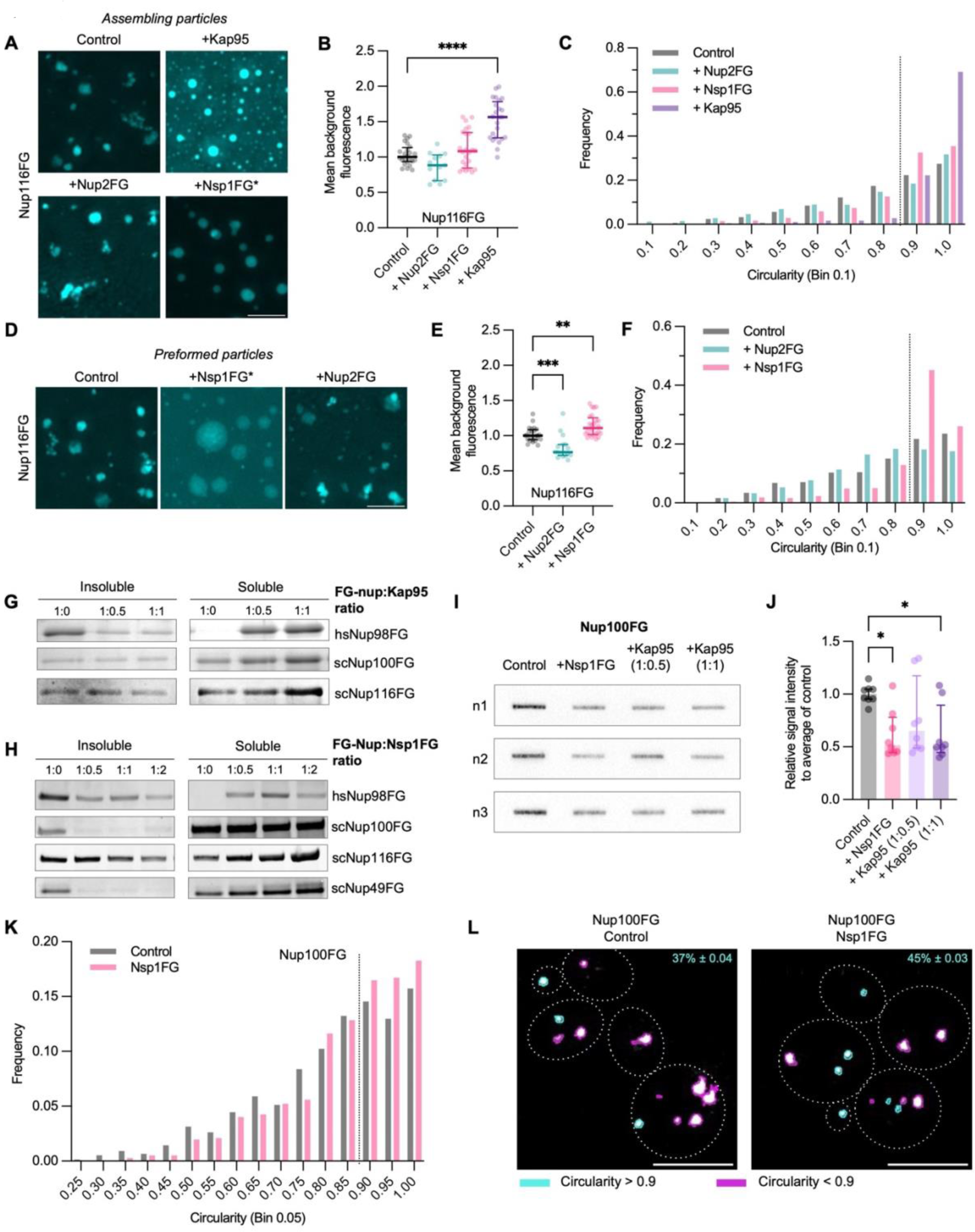
Nsp1 modulates FG-Nup phase state *in vitro* and in cells. (A) Representative z-projections of Nup116FG-FITC particles formed (1h) in the presence or absence of Kap95, Nsp1FG or Nup2FG. *Nsp1FG condition showed very heterogeneous particles (see Figure S4A). Scale bar, 5µm. (B) Mean background fluorescence of Nup116FG-FITC particles exemplified in (A). Graph shows median ± interquartile range of 24 images per condition (n=3). (C) Frequency distribution of the circularity of Nup116FG-FITC particles exemplified in (A). Bin width 0.1. 1000 particles per condition (n=3). (D) Representative images of preformed Nup116FG-FITC particles (1h) after addition of Nsp1FG or Nup2FG for another hour. *Nsp1FG condition showed very heterogeneous particles (see Figure S4J). Scale bar, 5µm. (E) Mean background fluorescence of particles exemplified in (D). Graph shows median ± interquartile range of 24 images per condition (n=3). (F) Frequency distribution of the circularity of particles exemplified in (D). Bin width 0.1. 1000 particles per condition (n=3). (G) Sedimentation assay of FG domains [3µM] with Kap95 [0, 1.5, 3 µM]. (H) Sedimentation assay of FG domains [3µM] with indicated concentrations of Nsp1FG [0, 1.5, 3, 6 µM]. (I) Filter trap assay of Nup100FG with indicated molar ratios of Nsp1FG and Kap95. (J) Quantification of filter trap assay. Graph shows median ± interquartile range (n=4 with 2 technical replicates per experiment). (K) Frequency distribution of the circularity of *in vivo* eGFP-Nup100FG particles in presence of absence of Nsp1FG. Bin width 0.05. >700 particles per condition (n=3). (L) Representative images of exemplified in (K). Percentages indicate the percentage particles with a circularity>0.9; mean ± SEM. Scale bar, 5µm. *p < 0.05, **p < 0.01, *** p < 0.001, ****p < 0.0001

A quantitative analysis of the particle size, fluorescence intensity and circularity, was performed using an image analysis pipeline that was validated to faithfully report on the phase state of FG-Nups (Bergsma et al. in preparation). Here, circularity can be used as a measure for the liquidity of condensates, where perfect circularity (a circularity of 1) aligns with the expected spherical morphology of a liquid droplet^18^, and the increase in background fluorescence can be used as a measure to detect the release of monomers into the solution. The parameters of particle fluorescence intensity and size cannot directly report on specific phase states but are able to report on particle changes in response to varying conditions. Quantitative analysis of 1000 particles per condition reports that in the presence of equimolar amounts of Kap95 a larger pool of soluble Nup116FG monomers exists and that the particles become rounder (Figure 3BC, Figure S4B). Additionally, there is a change in mean fluorescence of the particles (Figure S4D). This shows that Kap95 influences the phase state of the Nup116FG particles, i.e., stabilizing them in a more liquid-like state.

Next, we studied the effect of Nsp1FG (Nsp1_1-601_) on other FG-Nups and compared these effects to those of Kap95 and Nup2FG (Nup2_67-583_). Nup2FG has a similar length as Nsp1FG and its abundance also strongly decreases with aging, but its amino acid sequence lacks the bimodal charge distribution found in Nsp1. We observed that in the presence of Nsp1FG, the Nup116FG particles are very heterogeneous and assume circular shapes of various sizes and intensities (Figure 3A, and Figure S4A illustrates the variability). The quantitative assessment of the particles supports that they change and become more liquid-like when formed in the presence of equimolar ratios of Nsp1FG (i.e., significantly increased circularity (Figure 3C, Figure S4B) and significant changes in intensity and size (Figure S4CD)), while the addition of Nup2FG has no significant effects on circularity and background fluorescence (Figure 3A-C, Figure S4B-D). These effects of Nsp1FG are not readily detected on Nup49FG particles, possibly because they are already more liquid-like, as evidenced by their high circularity (Figure S4B-F).

In addition to the capacity of Nsp1FG to alter the FG-Nup phase state during particle assembly, we asked whether Nsp1FG also affects preformed Nup116FG particles. Adding Nsp1FG to preformed particles resulted in a heterogeneous population of particles (Figure 3D, Figure S4J). Also here, the quantitative analysis predicts that Nsp1 makes preformed Nup116FG particles more liquid-like (significantly increased circularity and background (Figure 3EF, Figure S4G) and significant changes in intensity and size (Figure S4HI)). All of this is not observed when adding Nup2FG, where rather a decrease in circularity and background were observed (Figure 3EF, Figure S4G-I).

To complement the imaging-based analysis of the phase state modulating properties of Kap95 and Nsp1FG on FG-Nups, we performed biochemical fractionation assays. Sedimentation assays show which fraction of the protein is in solution and which is in particles, be it in condensates or aggregates. Consistent with the imaging data, the presence of Kap95 decreases the insoluble fraction of the cohesive human Nup98FG (Nup98_50-547_) and mildly decreases the insoluble fraction of Nup116FG and Nup100FG (Figure 3G, Figure S4K). The presence of Nsp1FG also decreases the insoluble fractions of Nup98FG, Nup116FG, Nup100FG and Nup49FG (Figure 3H, Figure S4L). The effects of Kap95 and Nsp1FG are similar as observed with the addition of 1,6-hexanediol (Figure S4MN). Depending on the experimental conditions (e.g. addition of crowding agents, or a prolonged time frame for particle formation), a fraction of the Nup100FG particles can transition into SDS-insoluble aggregates that can be trapped in filter trap assays^54^. Interestingly, an anti-aggregation effect of Nsp1FG and Kap95 on Nup100FG is observed in filter trap assays (Figure 3IJ). The anti-aggregation effect of Kap95 had previously also been shown for the FG-domain of human Nup153^40^.

Altogether the imaging and biochemical analyses shows that both Nsp1FG and Kap95 promote a more liquid-like state of *in vitro* formed NupFG condensates and that they prevent or delay the formation of SDS-insoluble aggregates.

To evaluate whether the *in vitro* observations are also present in a cellular environment, we co-overexpressed the Nsp1FG domain with GFP-Nup100FG or GFP-Nup116FG in *S. cerevisiae*. As also shown in the accompanying paper by Kozai *et al.* and as reported previously^44, 47^ GFP-Nup100FG and GFP-Nup116FG form foci in cells while Nsp1FG localizes dispersed throughout the cytoplasm. A quantitative analysis of the effects of co-overexpressing Nsp1 indicates that the particles are more liquid-like (significantly increased circularity (Figure 3KL, Figure S5A) and significant changes in intensity (Figure S5C, no change in particle size (Figure S5B)). The frequency distribution of the particle circularity shows that Nup100FG and Nup116FG particles with a circularity below 0.9 are less prevalent when co-expressing Nsp1FG, while those above 0.9 are more frequent compared to the control (Figure 3KL, Figure S5D); this distribution is also observed in the *in vitro* experiments (Figure 3CF).

Taken together, these results show that Kap95 and Nsp1FG alter the phase state of other FG-Nups *in vitro* and likely also *in vivo*. The function of these proteins in the native Nsp1 condensate may thus be to keep FG-Nups in an assembly competent (phase) state.

### Dynamic and charged linker sequences of Nsp1 cover FG-Nup condensates

In order to gain a molecular understanding of the biochemical behavior of Nsp1 towards other FG-Nups, we performed coarse-grained molecular dynamics (CGMD) simulations using the 1BPA model for FG-Nups^77–79^. This model has been used previously to study nuclear transport^78, 80–84^ and to characterize the phase separation behavior of yeast FG-Nups^79^. We first simulated phase separation of Nup116FG, after which we added Nsp1FG monomers to the phase separated Nup116FG system. Similar to observations of previous simulations^79^, Nup116FG phase separates and forms a single condensate by the many dynamic FG–FG interactions, encompassing almost all protein chains (Figure S6A). By adding Nsp1FG monomers we observe that Nsp1FG monomers join the dense phase of Nup116FG. We find that only the head domains of Nsp1FG engage with the Nup116FG condensate, while the extended domains of Nsp1FG are predominantly pointing outward (Figure 4AB). To quantify the molecular interactions between Nup116FG and Nsp1FG, we calculated the contact frequencies (Figure 4C). The bimodal behavior of Nsp1FG towards Nup116FG is clearly visible, where almost all interactions are with Nsp1’s collapsed head domain, and very few with its extended domain. The contact map highlights that most interactions between Nup116FG and Nsp1FG are between FG-motifs. These motifs act as highly-dynamic hydrophobic stickers forming FG-FG contacts with life times as low as ∼0.5 ns^79^. As FG-motifs are equally abundant in both domains of Nsp1, we conclude that the bimodal behavior is not determined by the FG-motifs, but rather by the different nature of the spacers that are interspersed between the FG-motifs in the two domains.

**Figure 4.**
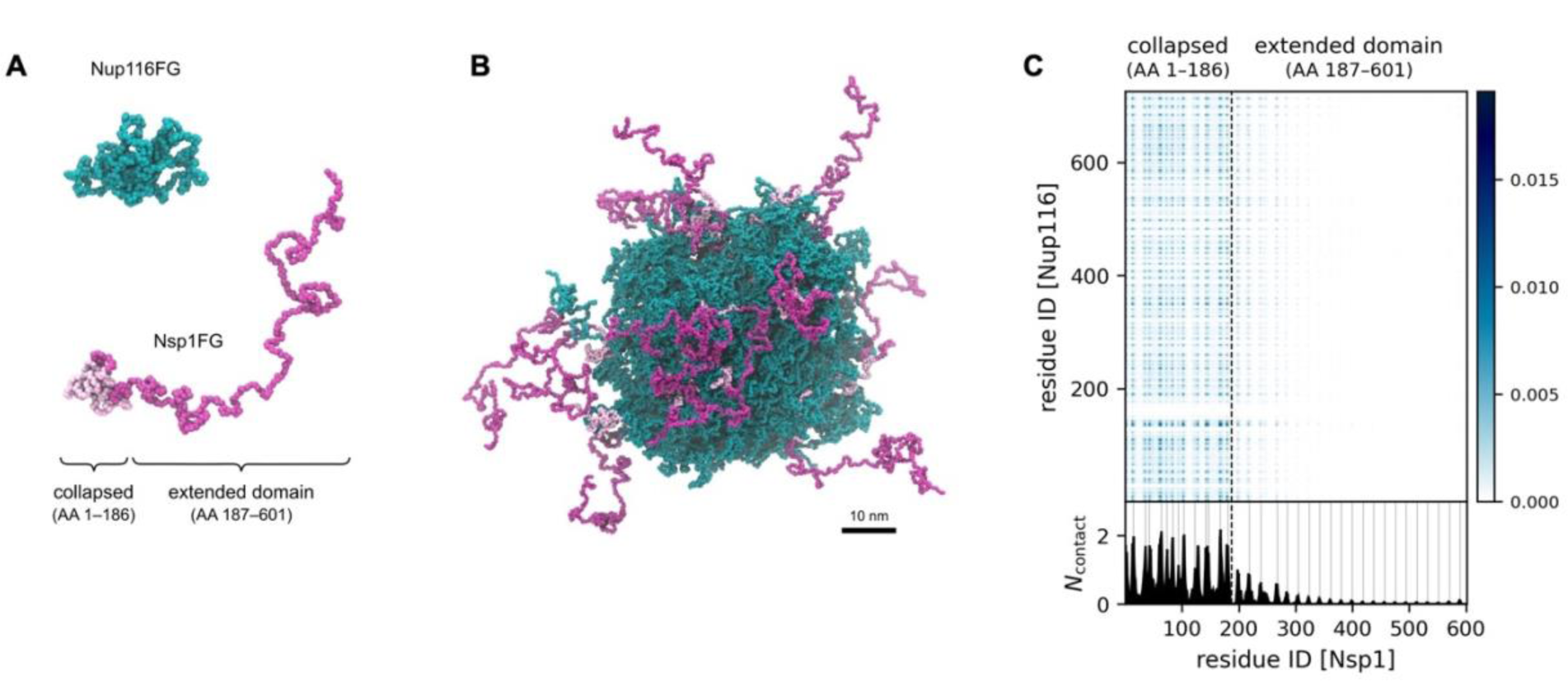
Coarse-grained modeling of phase separation of Nup116FG and Nsp1FG. (A) Snapshots of single chain conformations of Nup116FG and Nsp1FG, highlighting the bimodal character of Nsp1FG. (B) The cohesive collapsed domains (light pink) mainly interact with the Nup116FG condensate, while the extended domains (magenta) are mostly exposed to the solvent. Only Nsp1FG molecules that are in contact with the Nup116FG condensate are shown. (C) Average number (top) and one-dimensional summation (bottom) of intermolecular contacts per protein replica as a function of residue number. Gray lines indicate the location of the FG-motifs; black dashed line the boundary between the collapsed and extended domain.

The CGMD simulations thus suggest that the activity of Nsp1 towards maintaining a liquid-like behavior of Nup116 condensates *in vitro* is related to the presence of dynamic and charged linker sequences in the extended domain, which promotes their projection from the condensate. We propose that the bimodal behavior of Nsp1FG towards other FG-Nups mechanistically explains how Nsp1 can keep them in a more soluble or liquid-like state.

### Reducing Nsp1 expression in young cells mimics aging phenotypes

Having established that Nsp1 plays an important role in maintaining a liquid-like state of FG-Nup condensates, we investigated the cellular effects of reducing Nsp1 levels in young cells. For this purpose, a Nsp1 heterozygous diploid yeast strain was created (NSP1/nsp1). Mass spectrometry showed that Nsp1 levels are reduced to about 30% in these cells, which corresponds to Nsp1 levels reached in mid-aged cells (Figure 5A). Interestingly, the reduction of Nsp1 levels lead to a reduction in the cellular abundance of other FG-Nups (especially Nup2 and Nup60) but not of the scaffold nups (Figure 5B). This reduction of FG-Nups abundances was also observed in previous aging proteomics and in microscopy data on aged cells^21–23^, and suggests that a reduction of Nsp1 levels may be causal to the decreased FG-Nup levels in aging.

**Figure 5.**
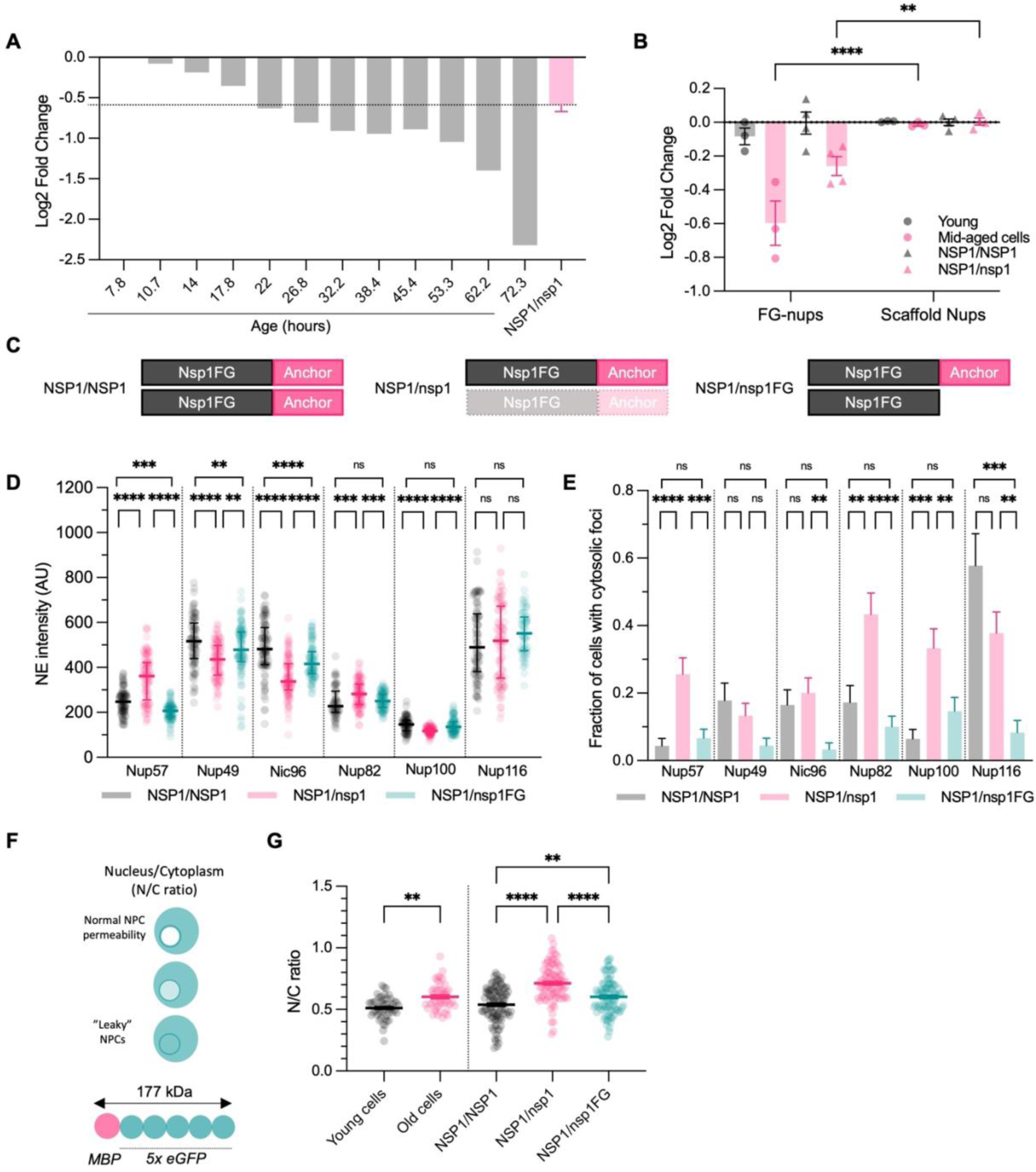
Reducing Nsp1 levels in young cells mimics aging phenotypes. (A) Reduction in Nsp1 protein levels in aging yeast cells^21^ compared to Nsp1 protein levels in NSP1/nsp1. Fold change is compared to t7.8 for the aging dataset and to NSP1/NSP1 for NSP1/nsp1. Graph shows mean ± SEM for NSP1/nsp1 (n=5). (B) Reduction in Nsp1 results in loss of FG-Nups, not scaffold nups. Graph shows mean ± SEM (n=3-5). (C) Graphical representation of NSP1/NSP1 (control), NSP1/nsp1 (Nsp1 heterozygote) and NSP1/nsp1FG (reintroduction Nsp1FG-domain) background strains. (D) Nuclear envelope (NE) intensity of eGFP tagged nups in the background strains of (C). Graph shows median ± interquartile range of >60 cells per condition (n=2-3). (E) Fraction of cells with cytosolic foci of eGFP tagged nups in the background strains of (C). Graph shows mean ± SEM of >60 cells per condition (n=2-3). (F) Graphical representation of the transport assay^80^. (G) N/C ratio of young and 20 hours batch-aged wild type cells and of control and Nsp1 mutant strains illustrated in (C). ns: not significant, **p < 0.01, *** p < 0.001, ****p < 0.0001

To investigate the effects of reduced Nsp1 levels on the NPC components more closely, the localization and levels of several endogenously tagged Nup-GFP fusions (Nup57, Nup49, Nup82, Nic96, Nup100 and Nup116) were determined by imaging. Clear alterations in fluorescence levels at the NE were observed, where some nup levels were increased (Nup57 and Nup82), whereas others showed reduced (Nup49, Nic96, Nup100) or unchanged (Nup116) protein levels (Figure 5D, see Figure S7B for whole cell levels). This is a clear indication that lowering the Nsp1 levels leads to a loss of stoichiometry of the NPC subunits, which, again, mimics what is observed in aging^21, 22^. The loss of Nup stoichiometry and NPC assembly problems in ageing occur post transcriptionally since Nup mRNA levels do not change as much in aging^21, 22^. Consistently, for Nup57, Nup82 and Nup100 there was an increase in cytosolic foci in mother cells, indicative of problems with their assembly into NPCs (Figure 5E). As our biochemical studies showed that the Nsp1FG domain has phase state modulating properties, which we hypothesized to have a surveillance role, we tested if introduction of just Nsp1FG may revert the phenotypes resulting from the reduction of the Nsp1 protein levels. Notably, Nsp1FG lacks the anchor domain and does not assemble into NPCs (Figure 5C, Figure S7A). Strikingly, reintroducing Nsp1FG rescues the loss of stoichiometry at the NPC, and it reduces the number of cytosolic foci (Figure 5DE). As a control, such effects are not observed with a Nup49 heterozygote (Figure S7CD). Clearly, the presence of the non-NPC resident pool of Nsp1FG is sufficient to revert the defects that are the result of reducing full length Nsp1 levels. This, together with the biochemical data, shows a surveillance function of non-NPC resident Nsp1FG for other FG-Nups, and likely also the native non-NPC resident pool of full length Nsp1 has this function.

To assess if the surveillance function of Nsp1 is also important for NPC functionality, the passive permeability of NPCs was determined. Under normal physiological conditions a large reporter containing a maltose binding domain (MBP) fused with five GFP proteins (MG5) is excluded from the nucleus, leading to a low ratio between the fluorescence in the nucleus over that in the cytosol (N/C ratio)^80^ (Figure 5F). In batch-aged cells the N/C ratio mildly increases, indicative of a reduced diffusion barrier in mid-aged cells (20h). When Nsp1 protein levels are reduced in young cells, as is the case in the NSP1/nsp1 heterozygous diploid, also an increase in N/C ratio is observed (Figure 5G). And indeed, the functional deficits of the NPC can be partially rescued by reintroducing only the Nsp1FG domain (Figure 5G) As will become clear below, this rescue is related to the cytosolic chaperone function of Nsp1 as Nsp1FG is not actually incorporated in NPCs. In the Nup49 heterozygote, also a small increase in N/C ratio of MG5 was observed, however not to the extent of the Nsp1 heterozygote (Figure S7E).

Together these results show that reduced Nsp1 levels in young cells phenocopies mid-aged cells, *i.e.* the phenotypes of decreased FG-Nup levels, loss of stoichiometry and transport defects, which were previously attributed to NPC assembly defects^22^. All these phenotypes can be rescued by reintroducing just the Nsp1FG-domain. We thus predict that reduced levels of Nsp1, and particularly of the FG-domain, may be causal to these NPC phenotypes in aged cells.

### Nsp1 intersects with classical chaperones of the protein quality control system

From the above we conclude that Nsp1 and especially its FG-domain alters the phase state of FG-Nups, similar to how molecular chaperones guard the stability of IDPs and FG-Nups in particular^54, 55, 85^. From the mass spectrometry data on NSP1/nsp1 cells we see that when Nsp1 protein levels are reduced in young cells, several heat shock proteins (Hsp104, Hsp42, Hsp78, Hsp82 and Ssa3/4) are upregulated (Figure 6A). This heat shock profile is very similar in aged yeast cells^21^ (Figure 6B). Strikingly, the heat shock response is normalized or even reduced when reintroducing only Nsp1FG (Figure 6AB). To further substantiate the relationship between Nsp1 and the protein quality control system, we expressed a protein stress reporter^86, 87^ in yeast cells. Under normal conditions cells are able to keep this unstable Luciferase-eGFP protein soluble, however when they experience protein stress, luciferase aggregates are formed. Indeed, in the NSP1/nsp1 strain we see an increase in the prevalence of Luciferase-eGFP aggregates, which is not observed in NUP49/nup49 and NUP116/nup116 strains (Figure 6C). The luciferase is stabilized when reintroducing the Nsp1FG domain (Figure 6C), further substantiating the surveillance function of the FG-domain of Nsp1.

**Figure 6.**
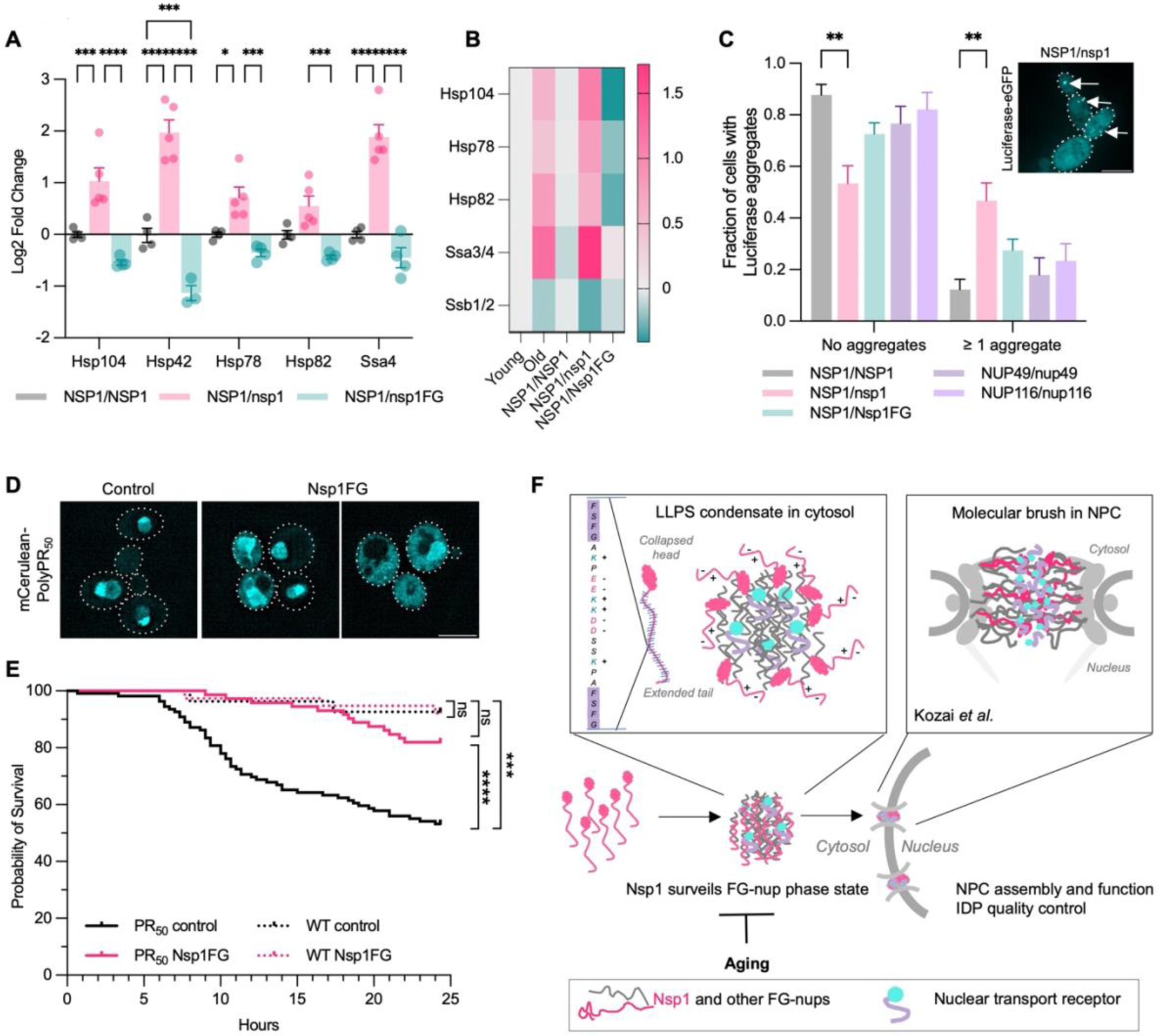
Nsp1 and protein quality control system. (A) Heat shock protein levels in control and Nsp1 mutant strains illustrated in (C). (n=5) (B) Heatmap of heat shock protein levels in control and Nsp1 mutant strains illustrated in (C) compared to aged yeast cells^21^. (C) Quantification of prevalence of Luciferase aggregates in indicated strain backgrounds. Graph shows mean ± SEM of 90 cells per condition (n=3). Inset: representative images of aggregates assay. White arrows indicate luciferase aggregates. Scale bar, 5µm. (D) Representative images of cells expressing mCerulean-PR50 alone (control) or together with Nsp1FG. Scale bar, 5µm. (E) 24-hour probability of survival of yeast cells expressing polyPR50 alone or together with Nsp1FG in presence or absence of Nsp1FG from time lapse imaging in microfluidic chambers. Controls are wildtype cells and cells expressing Nsp1FG. (F) Graphical representation of surveillance role of Nsp1 for FG-Nups ns: not significant, *p < 0.05, ** p < 0.001, *** p < 0.001, ****p < 0.0001

Since reducing Nsp1 levels leads to a more generalized protein stress response, we hypothesized that the chaperone-like effect of Nsp1 might be extendable to other IDPs. Since there is evidence that Nup62, the human orthologue of Nsp1, is linked to ALS and FTD related pathology^88–90^, we tested the effect of Nsp1FG expression on toxicity and localization of polyPR, a peptide linked to C9orf72 ALS. In yeast cells, as in many other models, mCerulean-tagged polyPR localizes in the nucleolus and nucleus and its expression is toxic^30, 91^. Previous work connected the toxicity of polyPR with its localization in the nucleolus^92–96^. Overexpression of Nsp1FG together with mCerulean-polyPR changes the subcellular localization of mCerulean-polyPR (Figure 6D, Figure S8D). When overexpressing Nsp1FG in cells expressing untagged polyPR, the toxicity is mildly reduced (Figure S8A). Following single cells in a microfluidic chip^66^ for 24h provides a more precise measure of cell survival. The expression of polyPR slowed down cell division and drastically reduced survival to ∼53% after 24h. The controls, wildtype cells or wildtype cells overexpressing Nsp1FG, both have normal division times and a survival of ∼92% (Figure 6E, Figure S8BC). When polyPR is expressed together with Nsp1FG, we observe an impressive increase in the 24h cell survival from 53% to ∼82%. This rescue is not due to reduced polyPR expression levels (Figure S8EF), hence we conclude that Nsp1FG attenuates the toxicity of polyPR.

Together these results show that reducing Nsp1 levels creates protein stress, and that the FG-domain of Nsp1 can alleviate this, as is evident from its ability to restore or even reduce normal chaperone expression levels, reduce luciferase aggregates, and counter polyPR toxicity.

## Discussion

In this study we addressed how the reduced expression of Nsp1 in aged cells relates to age-related NPC assembly problems. Especially in a protein complex as complicated as the NPC, a proper quality control system is crucial. We discovered that Nsp1 and Kap95 act as phase state modulators keeping FG-Nup condensates in a liquid-like state. We propose that this activity is needed to safeguard the FG-Nups while they are stored in a transient daughter cell specific condensate of FG-Nups and NTRs. The pool of Nsp1 that is not embedded in NPCs thus has a surveillance/chaperone function for IDPs and it shares this function with NTRs^32, 40^ and chaperones^54, 55, 97^ (Figure 6F).

### From a cytosolic condensate to a brush conformation inside the NPC

Our findings highlight that Nsp1 has an important additional role outside the central channel of the NPC. In addition to the NPC resident pool and a daughter specific focus^63^, we demonstrate that Nsp1 has a significant cytosolic pool. We show that the daughter specific focus is a condensate containing several other FG-Nups and the NTR Kap95. This condensate was already linked to NPC inheritance and assembly^63, 64^, and we extend on that by hypothesizing it serves as an assembly platform for new NPCs, in which Kap95 and Nsp1 keep FG-Nups in an assembly competent state until incorporation. Nup condensates with a function as an assembly platform have been previously described in *Drosophila* oogenesis^56^ and HeLa cells^57^. This function of nup condensates aligns with that of other condensates also aiding in assembly (e.g. ribosomes in the nucleolus) or acting as storage compartments (e.g. stress granules)^17, 19, 98^. In the accompanying paper, Kozai *et al.* show that once assembled into the NPC, the FG-Nups form a dynamic brush. This metamorphosis from a condensate state to a brush state, each having vastly different properties and functions, exemplifies the unique structural and functional versatility that IDPs have. Future research may reveal how the condensates resolve during NPC assembly and how the transition from condensate to brush is exactly controlled.

### Nsp1 maintains native FG-Nup condensates in a liquid-like state

In several biochemical *in vitro* and *in vivo* assays, we observe that Nsp1 alters the phase state of FG-Nups towards more liquid-like and soluble states and that Nsp1 delays formation of SDS-insoluble aggregates. Nsp1 has a striking amino acid sequence in its FG-domain. Like several other FG-Nups, Nsp1 is known to have a bimodal charge distribution along its sequence, resulting in a more collapsed head and an extended domain^47^ Unique to Nsp1 is that it has many FG repeats in the extended domain, and that the FG repeats are very evenly distributed and interspaced by linker sequences containing many positively and negatively charged amino acids (Figure S1C). Modeling predicts that Nsp1 is recruited to the condensates by FG-FG interactions with the collapsed head region, while the dynamic extended domain projects out of the condensate exposing the charged linker regions. We propose that the highly charged linker regions provide the condensates with surface properties that allow more rapid exchange of monomers leading to a more liquid-like condensate and to the release of monomers.

### Nsp1 surveillance within the NPC

The FG-Nups within the NPC do not form self-assembled condensates but are organized by the NPC’s scaffold to form a highly dynamic brush, as shown in the accompanying paper by Kozai *et al*. Interestingly, except for Nsp1, the FG-Nups in the equator of the central channel are mainly GLFG-Nups. In isolation GLFG nups have a stronger tendency to self-associate into condensates than other FG-Nups^41, 44, 47, 79^. This natural tendency encoded in the amino acid sequence is possibly actively inhibited within the NPC in several ways. The formation of stable FG-FG contacts is likely prevented by anchoring to the NPC scaffold, and by engagement with NTRs (this study and^32^) and chaperones^54, 55^. From the work presented here we can add Nsp1 to this list of possible guardians of FG-Nups within the NPC. Current models show that NPC assembly occurs through an in-side out mechanism^8, 99^ meaning that FG-Nups are imported into the nucleus prior to their assembly, an event that likely needs chaperoning of the FG-Nups. Many transcriptional regulators and RNA binding proteins also have large ID regions and many are prone to phase separate^17, 20^. We speculate that the chaperone function of Nsp1 may be particularly relevant when IDPs, including FG-Nups and transcriptional regulators, transit the NPCs.

### Surveillance of IDPs is where NPC biology and protein quality control intersect

Reducing Nsp1 levels in young cells results in several NPC assembly-related problems, such as loss of NPC stoichiometry and aggregation of several nups, in addition to problems with NPC transport function. Strikingly, these problems can be rescued by reintroducing just the Nsp1FG domain, which resides in the cytosol and is unable to anchor in the NPC. This indicates that cytosolic Nsp1FG exhibits a chaperone-like function to stabilize FG-Nups for proper incorporation into NPCs. In line, reducing Nsp1 levels also leads to upregulation of several heat shock proteins, which could be a compensatory mechanism for the chaperone-like activity of Nsp1. A strong indication is that when reintroducing just the Nsp1FG-domain, this stress response is again normalized, and protein stress is alleviated. The results with the polyPR expressing yeast show that the chaperone-like function of Nsp1 is not limited to FG-Nups but can be extended towards other IDPs. Our data contributes to a larger emerging theme of a tightly regulated interplay between FG-Nups, NTRs and the classical protein quality control system^54, 55, 85, 100, 101^. Interestingly, Nup62, the mammalian ortholog of Nsp1, has been found to participate in assemblies with disease-related IDPs, such as TDP43^89, 90^ and FUS^88, 102^. In general, protein quality control of stably folded proteins is much better understood^100, 101^ than that of IDPs. Aberrant phase state transitions of diverse IDPs occur in aged cells and in (models of) neurodegenerative diseases^73, 75, 76^, and a better understanding of what regulates these transitions is much needed. This study identifies Nsp1 as a new player in the field at the crossroads of NPC biology and protein quality control.

## Acknowledgments

We thank Annemiek Veldsink, Elizabeth Riquelme Barrientos and Joris van der Lienden for sharing reagents and Amarins Blaauwbroek and Leila Saba for technical support. We thank Dirk Görlich for the gift of NupFG constructs, Judith Frydman for the Luciferase construct and Anton Khmelinskii for the tandem fluorescent timer constructs. We thank Cor Calkhoven, Michael Chang, Michael Rout and all members of the Veenhoff and Chang laboratories for valuable suggestions.

This work was financially supported by the Netherlands Organization of Scientific Research grant no. VI.C.192.031 to LMV (T.A.O, L.M.V, A.S) and OCENW.GROOT.2019.068 (T.B., M.D., P.G.). T.A.O. is supported by a PhD fellowship from the Graduate School of Medical Sciences from the University Medical Center Groningen. This work made use of the Dutch national e-infrastructure with the support of the SURF Cooperative using grant no. EINF-3473.

## Author contributions

T.A.O and L.M.V conceived the project. T.A.O. and L.M.V wrote the manuscript with input of all authors. L.M.V., T.A.O. and P.R.O secured funding. A.S. provided expertise and feedback throughout the project. T.O. performed all experiments except those in Fig.3A-F,IJ and Fig. 4. T.B. performed microscopy-based condensate analysis of Fig.3A-F,IJ. M.D performed simulations in Fig.4 under supervision of P.R.O. J.C.W generated and analyzed proteome data, S.N.M. and P.G. provided reagents and expertise with microfluidics and *in vivo* phase separation assays, respectively.

---

**Figure S1.**
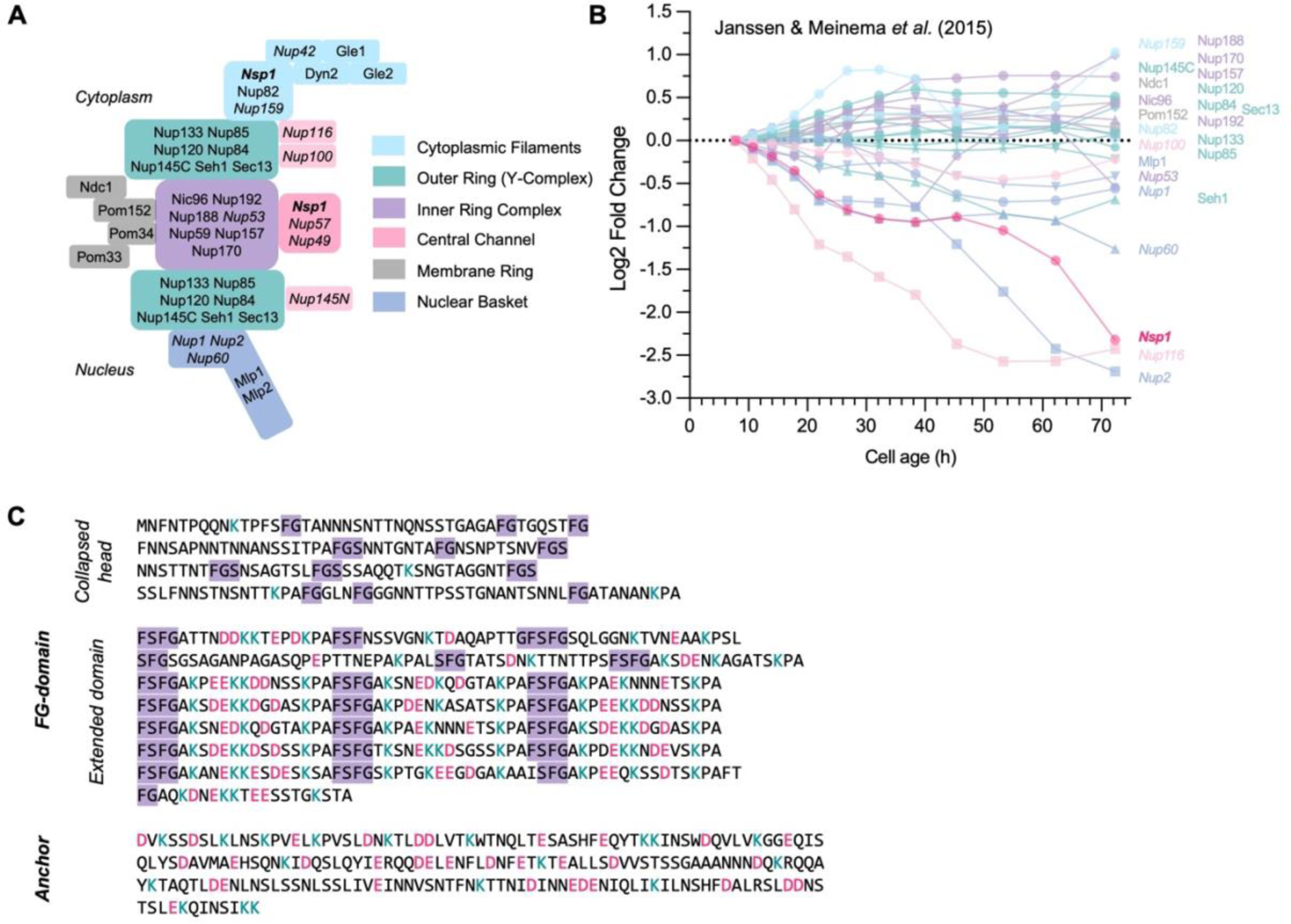
Nsp1 decreases with aging. (A) Detailed graphical representation of the orientation of the components in one spoke of the Nuclear Pore Complex. (B) Nup abundances in whole cell extracts of replicative aging cells normalized to young levels illustrated the loss of stoichiometry in aging. Data from Janssen & Meinema *et al.* (2015)^21^. (C) Amino acid sequence of Nsp1 with FG repeats highlighted and positively and negatively charged residues color coded in turquoise and magenta, respectively. Three distinct domains are indicated.

**Figure S2.**
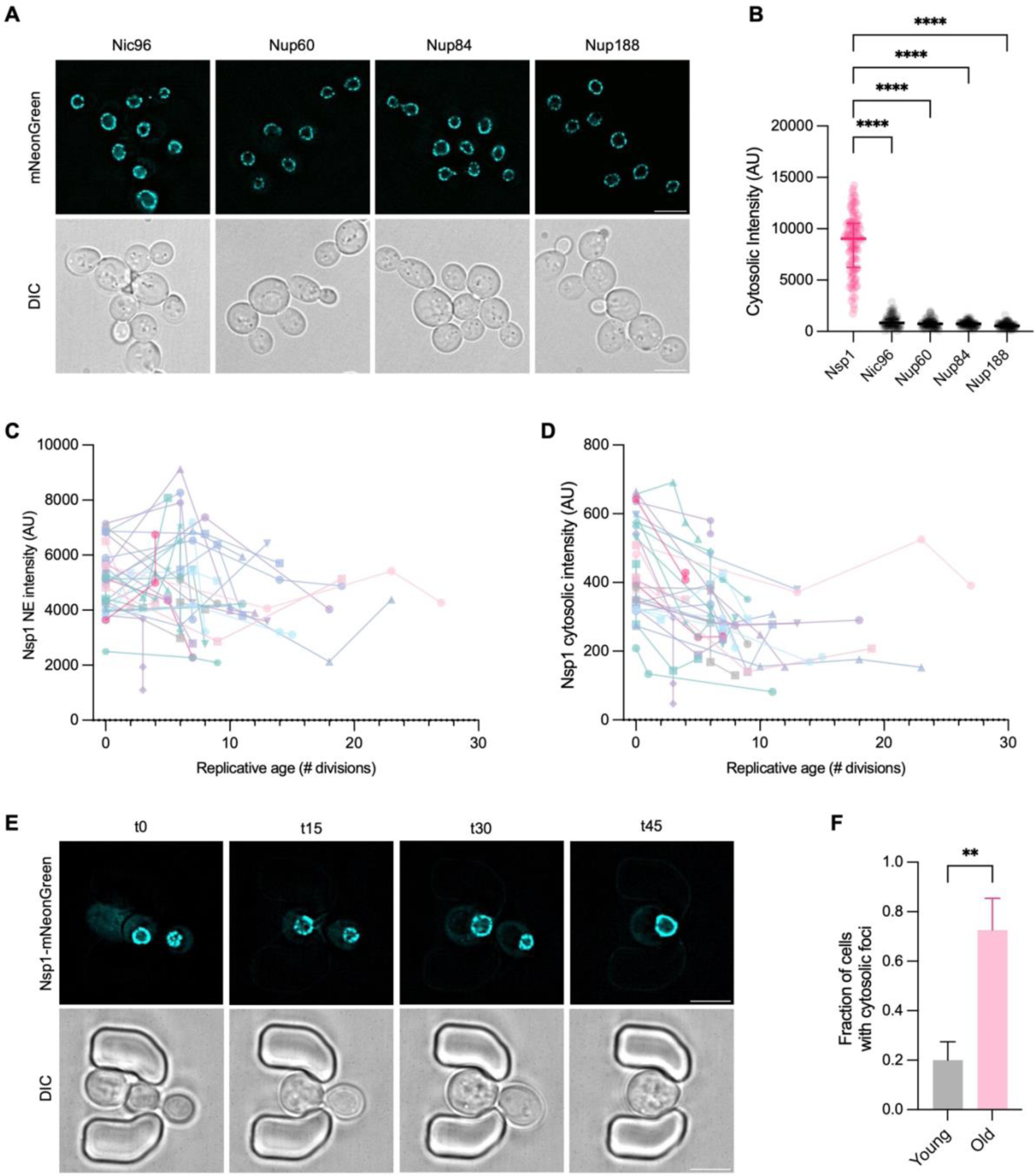
Nsp1 localizes in three distinct pools and decreases with aging. (A) Representative image of cells endogenously expressing mNeonGreen-tagged nups. Scale bar, 5µm. (B) Quantification of cytosolic levels of endogenously mNeongreen-tagged nups. Graph shows median ± interquartile range of 100 cells per condition. (C) Single cell trajectories of Nsp1 NE intensity levels at their corresponding replicative age. >30 cells (n=2). (D) Single cell trajectories of Nsp1 cytosolic intensity levels at their corresponding replicative age. >30 cells (n=2). (E) Representative image of Nsp1-mNeonGreen microfluidic timelapse. Scale bar, 5µm. (F) Fraction of cells with cytosolic foci in young and old cells. These foci are distinct from the daughter specific Nsp1 foci observed in young dividing mother cells and likely reflect aggregates. Graph shows mean ± SEM of >30 cells per condition. **p < 0.01, ****p < 0.0001

**Figure S3.**
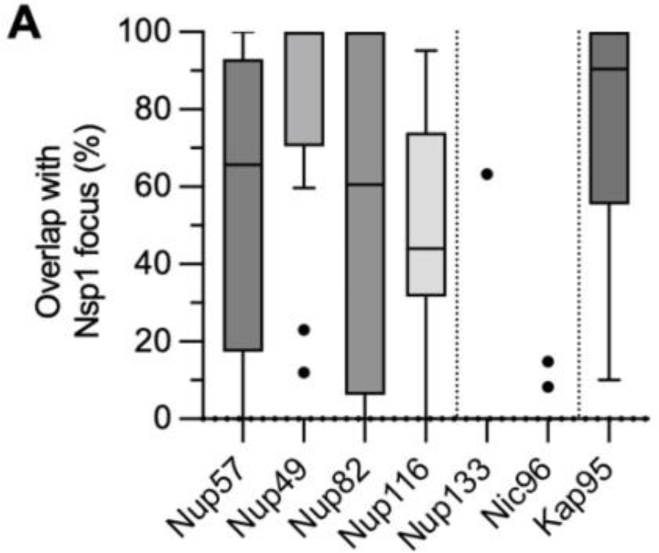
Characterization of native cytosolic Nsp1 condensate. (A) Percentage of overlap between signal from Nup-GFP and Nsp1-antibody in the Nsp1 focus in immunofluorescence. Graph shows boxplot.

**Figure S4.**
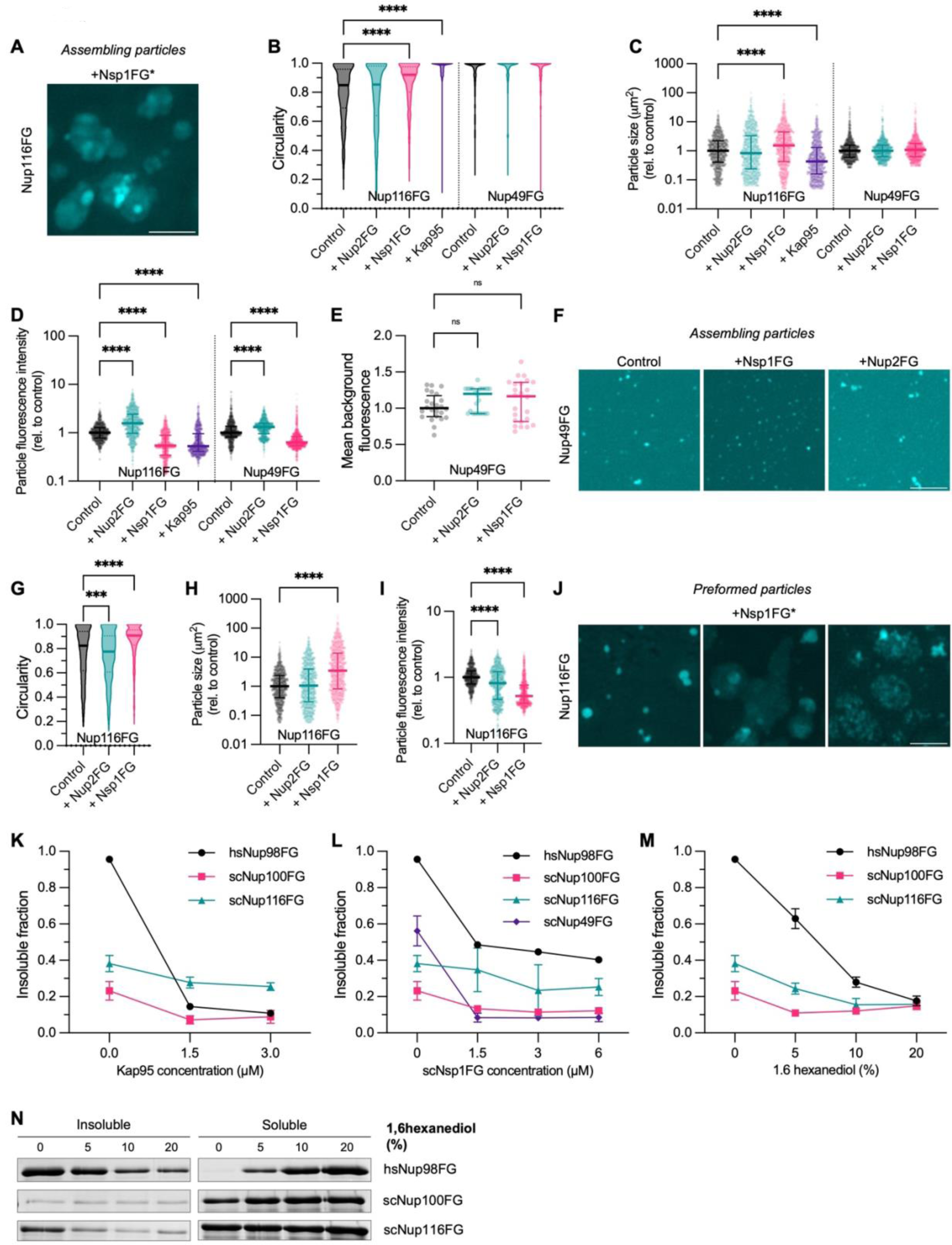
Nsp1 modulates FG-Nup phase state *in vitro*. (A) Representative images of the heterogeneity of assembling Nup116FG-FITC particles in the presence of Nsp1FG (1h). Scale bar, 5µm. (B) Circularity of assembling Nup116FG-FITC and Nup49FG-FITC particles. Graph shows median ± interquartile range of 1000 particles per condition (n=3). (C) Particle size of assembling Nup116FG-FITC and Nup49FG-FITC particles. Graph shows median ± interquartile range of 1000 particles per condition (n=3). (D) Particle fluorescence intensity of assembling Nup116FG-FITC and Nup49FG-FITC particles. Graph shows median ± interquartile range of 1000 particles per condition (n=3). (E) Mean background fluorescence of assembling Nup49FG-FITC particles. Graph shows median ± interquartile range of 24 images per condition (n=3). (F) Representative images of Nup49FG-FITC particles without or with Nsp1FG or Nup2FG. Scale bar, 5µm. (G) Circularity of preformed Nup116FG-FITC particles. Graph shows median ± interquartile range of 1000 particles per condition (n=3). (H) Particle size of preformed Nup116FG-FITC particles. Graph shows median ± interquartile range of 1000 particles per condition (n=3). (I) Particle fluorescence intensity of preformed Nup116FG-FITC particles. Graph shows median ± interquartile range of 1000 particles per condition (n=3). (J) Representative images of the heterogeneity of preformed Nup116FG-FITC particles after addition of Nsp1FG for 1h. Scale bar, 5µm. (K) Quantification sedimentation assay of FG domains [3µM] with Kap95. Graph shows mean ± SEM (n=3). (L) Quantification sedimentation assay of FG-Nups [3µM] with Nsp1FG. Graph shows mean ± SEM (n=3). (M) Quantification sedimentation assay of FG-Nups [3µM] with different concentrations of 1,6-hexanediol. Graph shows mean ± SEM (n=3). (N) Representative image of sedimentation assay of FG domains [3µM] with different concentrations of 1,6-hexanediol. **p < 0.01, ****p < 0.0001

**Figure S5.**
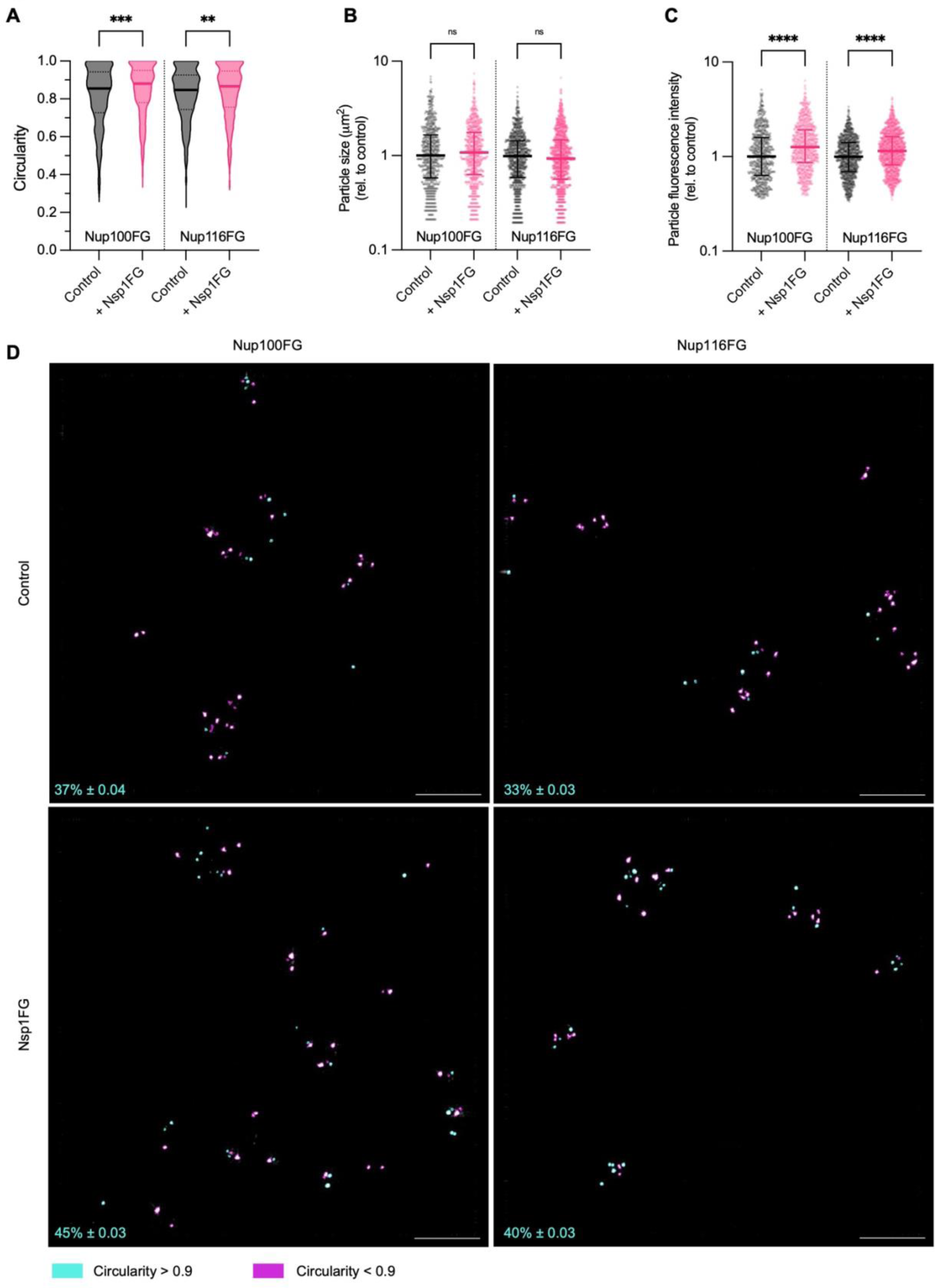
Nsp1 modulates FG-Nup phase state in cells. (A) Circularity of *in vivo* eGFP-Nup100FG and eGFP-Nup116FG particles in presence or absence of Nsp1FG. Graph shows median ± interquartile range of >700 particles per condition (n=3). (B) Particle size of *in vivo* eGFP-Nup100FG and eGFP-Nup116FG particles in presence or absence of Nsp1FG. Graph shows median ± interquartile range of >700 particles per condition (n=3). (C) Particle fluorescence intensity of *in vivo* eGFP-Nup100FG and eGFP-Nup116FG particles in presence or absence of Nsp1FG. Graph shows median ± interquartile range of >700 particles per condition (n=3). (D) Representative whole fields of view of frequency distribution of the circularity of *in vivo* eGFP-Nup100FG and eGFP-Nup116FG particles in presence or absence of Nsp1FG. Percentages show mean ± SEM. Scale bar, 5µm. ns: not significant, **p < 0.01, *** p < 0.001, ****p < 0.0001

**Figure S6.**
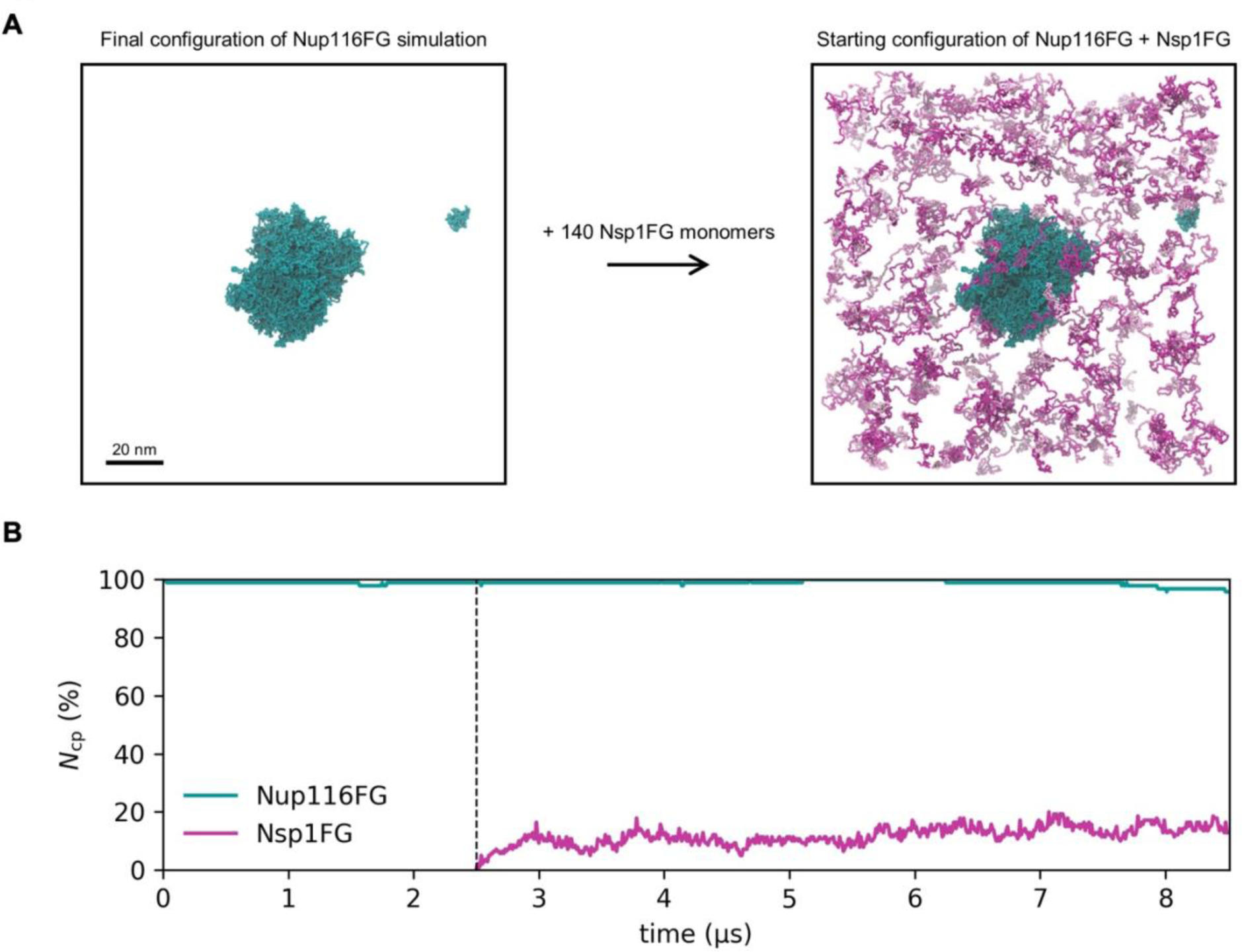
Coarse-grained modeling of phase separation of Nup116FG and Nsp1FG. (A) Starting configuration of the Nup116 with Nsp1 simulation. The final coordinates of the equilibration simulation of Nup116 are used as input, where 140 Nsp1 monomers are randomly placed in the simulation box, whilst avoiding any overlap with the monomers that are already present. (B) Fraction of FG-Nups that are in the condensate, Ncp (%), as a function of simulation time. The Nup116FG phase separation simulation started from a condensate structure, therefore Ncp = 100% at t^∗^ = 0. After 2.5 μs (marked by the dashed line) the Nsp1FG monomers are added to the simulation.

**Figure S7.**
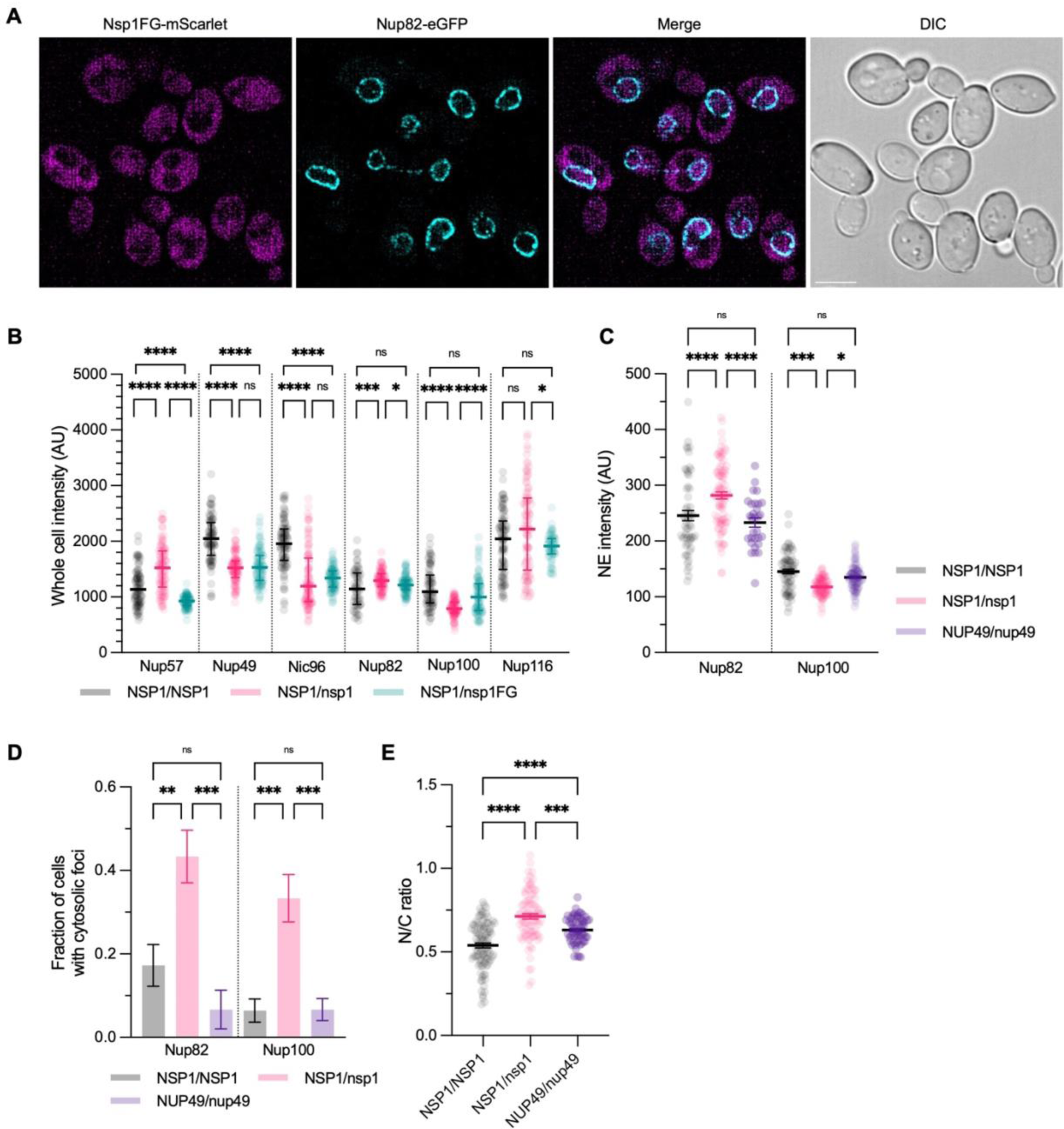
Reducing Nsp1 levels in young cells mimics aging phenotypes. (A) Representative images of cytosolic Nsp1FG-mScarlet localization in Nup82-eGFP strain. Scale bar, 5µm. (B) Whole cell intensity of eGFP tagged nups in background strains of Figure 5C. Graph shows median ± interquartile range of >60 cells per condition (n=2-3). (C) Nuclear envelope (NE) intensity of Nup82-eGFP and Nup100-eGFP in the NUP49/nup49 background. Graph shows median ± interquartile range of >60 cells per condition (n=2-3). (D) Fraction of cells with cytosolic foci of Nup82-eGFP and Nup100-eGFP in the NUP49/nup49 background. Graph shows median ± interquartile range of >60 cells per condition (n=2-3). (E) N/C ratio of NUP49/nup49 background control strain. Graph shows mean ± SEM of >60 cells per condition (n=3). ns: not significant, *p < 0.05, **p < 0.01, *** p < 0.001, ****p < 0.0001

**Figure S8.**
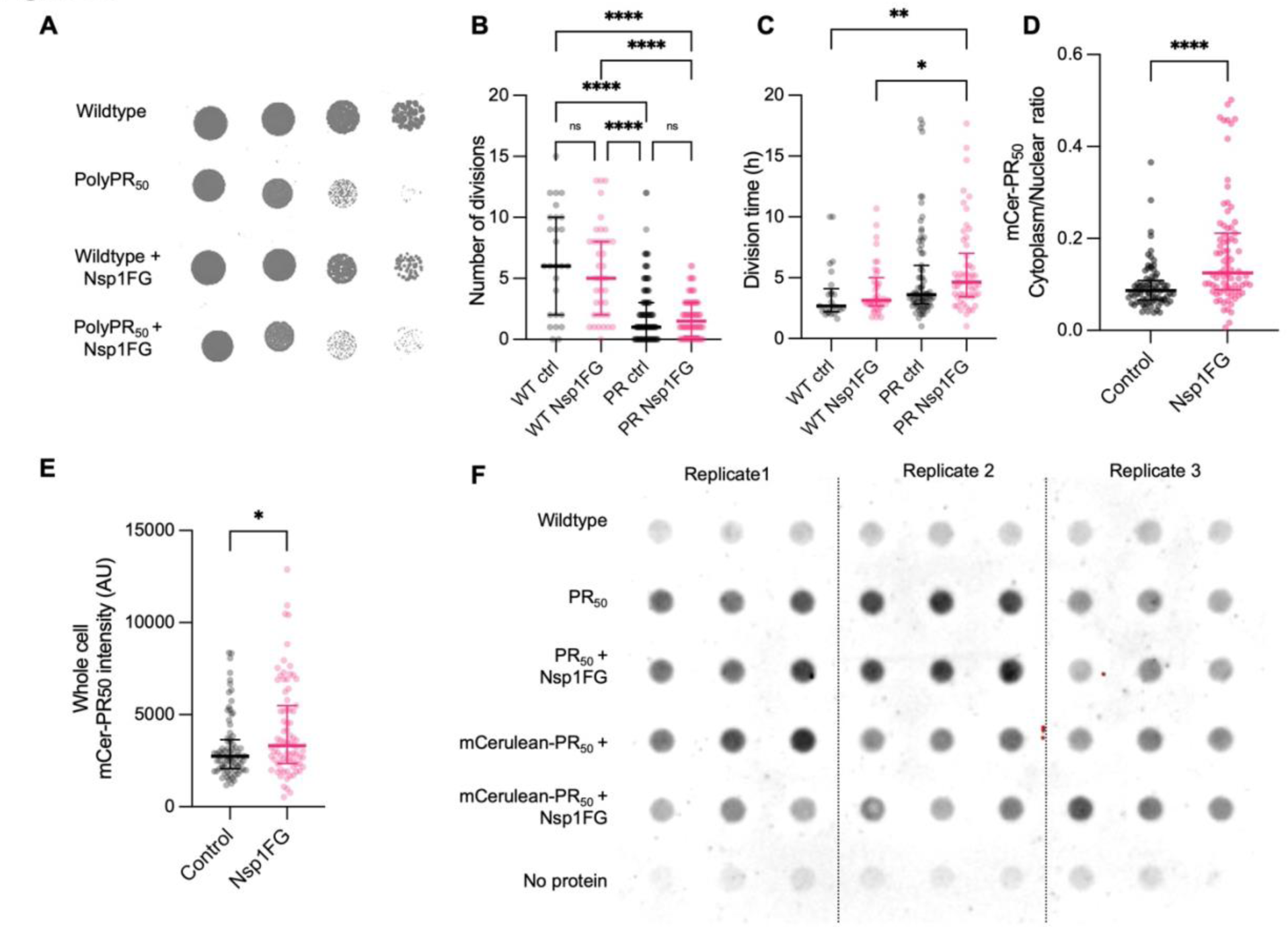
Nsp1 and protein quality control system. (A) Viability of yeast cells expressing polyPR50 alone or together with Nsp1FG. Controls are wildtype cells and cells expressing Nsp1FG. (B) Number of cell divisions of yeast cells expressing polyPR50 in presence or absence of Nsp1FG compared to wildtype cells. Graph shows median ± interquartile range of >27 cells per condition (n=2-3). (C) Division time of yeast cells expressing polyPR50 in presence or absence of Nsp1FG compared to wildtype cells. Graph shows median ± interquartile range of >27 cells per condition (n=2-3). (D) Quantification of mCerulean-PR50 signal in the cytoplasm and nucleus of cells exemplified in (Figure 6D). Graph shows median ± interquartile range of 80 cells per condition (n=2). (E) Whole cell mCerulean-PR50 intensity in absence or presence of Nsp1FG. Graph shows median ± interquartile range of 80 cells per condition (n=2). (F) Dot blot of yeast cells expressing polyPR50 or mCerulean-PR50 in presence or absence of Nsp1FG compared to wildtype cells (n=3). ns: not significant, *p < 0.05, **p < 0.01, ****p < 0.0001

## Materials and Methods

Key source table

| Reagents and Resources | Source | Identifier |
| --- | --- | --- |
| **Antibodies** |  |  |
| Monoclonal antibody mouse anti-GFP | Santa Cruz | sc-9996 |
| HRP-conjugated mouse IgG kappa binding protein | Santa Cruz | sc-516102 |
| mouse monoclonal (32D6) anti-Nsp1 | Abcam | ab4641 |
| anti -mouse-AF594F(ab')<sub>2</sub>-Goat anti-Mouse IgG (H+L) Cross-Adsorbed Secondary Antibody, Alexa Fluor™ 594 | Thermo Scientific | A-11020 |
| cMyc antibody (clone 9E10) | Takara Bio | 631206 |
| **Chemicals** |  |  |
| Drop out media amino acids | Sigma-Aldrich |  |
| D-Glucose anhydrous | Fisher Chemical™ | 10141520 |
| D-Raffinose pentahydrate | Thermo Scientific | 195675000 |
| D-Galactose | Acros Organics | 150610010 |
| Phosphate-buffered saline | Sigma-Aldrich | P4417 |
| Tris base | Fisher Scientific™ | BP152-1 |
| HEPES | Fisher Scientific™ | BP310-500 |
| Sodium chloride | Acros Organics | 207790010 |
| Tween20 | MP Biomedicals | TWEEN201 |
| Magnesium chloride hexahydrate | Sigma-Aldrich | M2393 |
| 1,4-Dithiothreitol (DTT) | Roche | 10708984001 |
| Glycerol | Sigma-Aldrich | G5516 |
| Phenylmethanesulfonyl fluoride (PMSF) | Sigma-Aldrich | P7626 |
| cOmplete ULTRA tablets, Mini EDTA-free | Roche | 05892791001 |
| Albumin bovine serum (BSA) | Acros Organics | 268131000 |
| Glass beads | BioSpec Products | 11079105 |
| Pierce™ BCA Protein Assay Kit | Fisher Scientific™ | 23225 |
| ECL Prime Western Blotting Detection Reagent | Amersham | RPN2232 |
| GX Stain-Free™ FastCast™ Acrylamide Kit, 10% | BioRad | 1610183 |
| sulfo-NHS-LC-biotin | Thermo scientific | 21335 |
| Streptavidin-coated magnetic beads | Brunschwig | 84660-5 |
| Fluorescent brightener 28 | Sigma-Aldrich | F3543 |
| Zymolyase T20 | Amsbio | 120491-1 |
| D-Sorbitol | Thermo-Scientific | 036404.A3 |
| Brilliant Blue G | Sigma-Aldrich | G-250 |
| fluorescein-5-maleimide | Thermo-Scientific | 62245 |
| NucBlue Live Cell stain | Invitrogen | R37605 |
| Sequencing grade modified trypsin | Promega | V5111 |
| Guanidine Hydrochloride (GuHCl) | Thermo Scientific | 24110 |
| Paraformaldehyde | Acros | A0319048 |
| Concanavalin A | Sigma-Aldrich | L7647 |
| Acetone | Sigma-Aldrich | 15A220519 |
| Nitrocellulose membrane Protran 0.2um | Amersham | GE10600001 |
| Ammonium persulfate | Merck | A3678 |
| NaOAc | Fisher Chemical™ | S2080/53 |
| TEMED | Fisher Scientific™ | 115664 |
| MgOAc | Fisher Scientific™ | BP215 |
| TritonX100 | Acros Organics | 215682500 |
| NuPAGE™ 4 to 12%, Bis-Tris, 1.0 mm, Midi Protein Gels | Invitrogen | WG1403BOX |
| EDTA | Sigma-Aldrich | ED2P-500 |
| Sodium chloride | Acros Organics | 207790010 |
| Methanol Technical | VWR | 20903.368 |
| IPTG | Sigma-Aldrich | 10724815001 |
| Ni sepharose | Cytiva | 17531802 |
| 1,6-hexandiol | Sigma-Aldrich | 240117-50 |
| **Software and Algorithms** |  |  |
| ImageJ | 103 | https://imagej.nih.gov/ij/ |
| softWoRx | Cytiva | n.a. |
| Prism 9 | Graphpad |  |

### Strains and growth conditions

All *Saccharomyces cerevisiae* strains used in this study had a BY4741 (haploid) or BY4743 (diploid) background. C-terminal tags and gene deletions were created using the yeast PCR toolbox^104^. All used strains are listed in the Stain table.

Yeast cells were grown in Synthetic drop-out (SD) medium supplemented with 2% (w/v) D-glucose at 30 °C, 200rpm. For strains with a galactose inducible plasmid; stains were first grown overnight in 2% D-glucose before switching to 2% (w/v) D-(+)-raffinose and induced with 0.1% (w/v) D-galactose for 1h (pPP008-MBP-5x eGFP) or 24h (pAG426-LucDM-eGFP) or with 1% (w/v) for 1h (pACM021-eGFP-Nup100FG, pACM021-eGFP-Nup116FG). Experiments were performed with cells in the exponential growth phase (OD∼0.5-0.8). Unless mentioned otherwise, 1mL of culture was concentrated (2400rcf, 2min) to 100uL of which 3uL was mounted on a glass slide with coverslip.

### Strain table

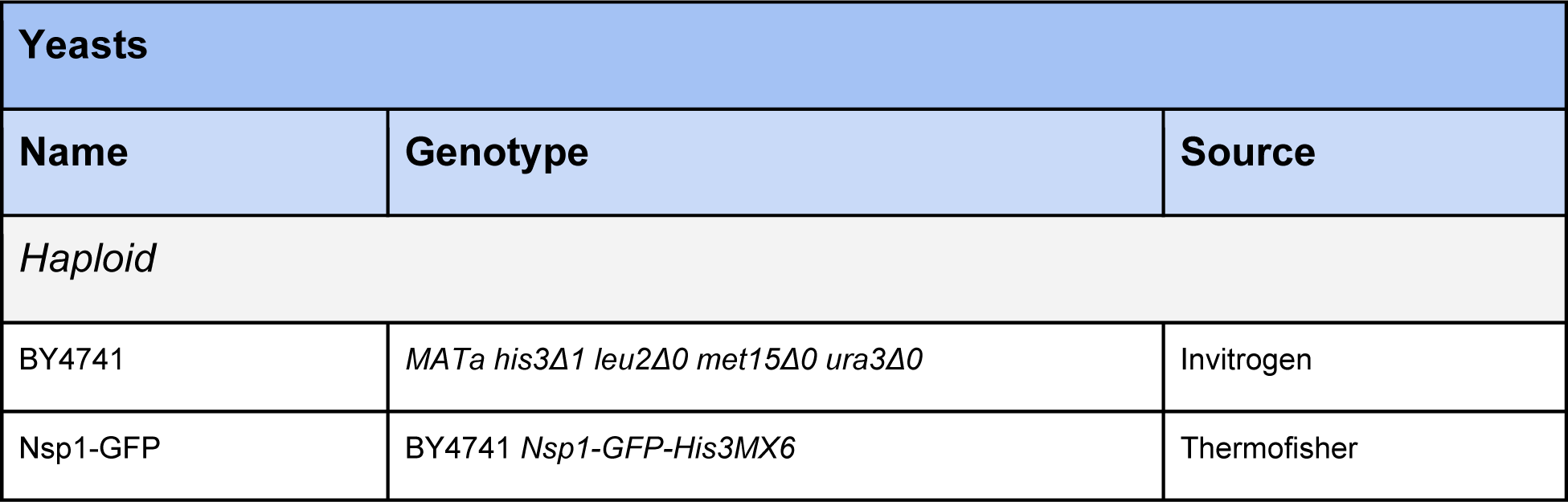
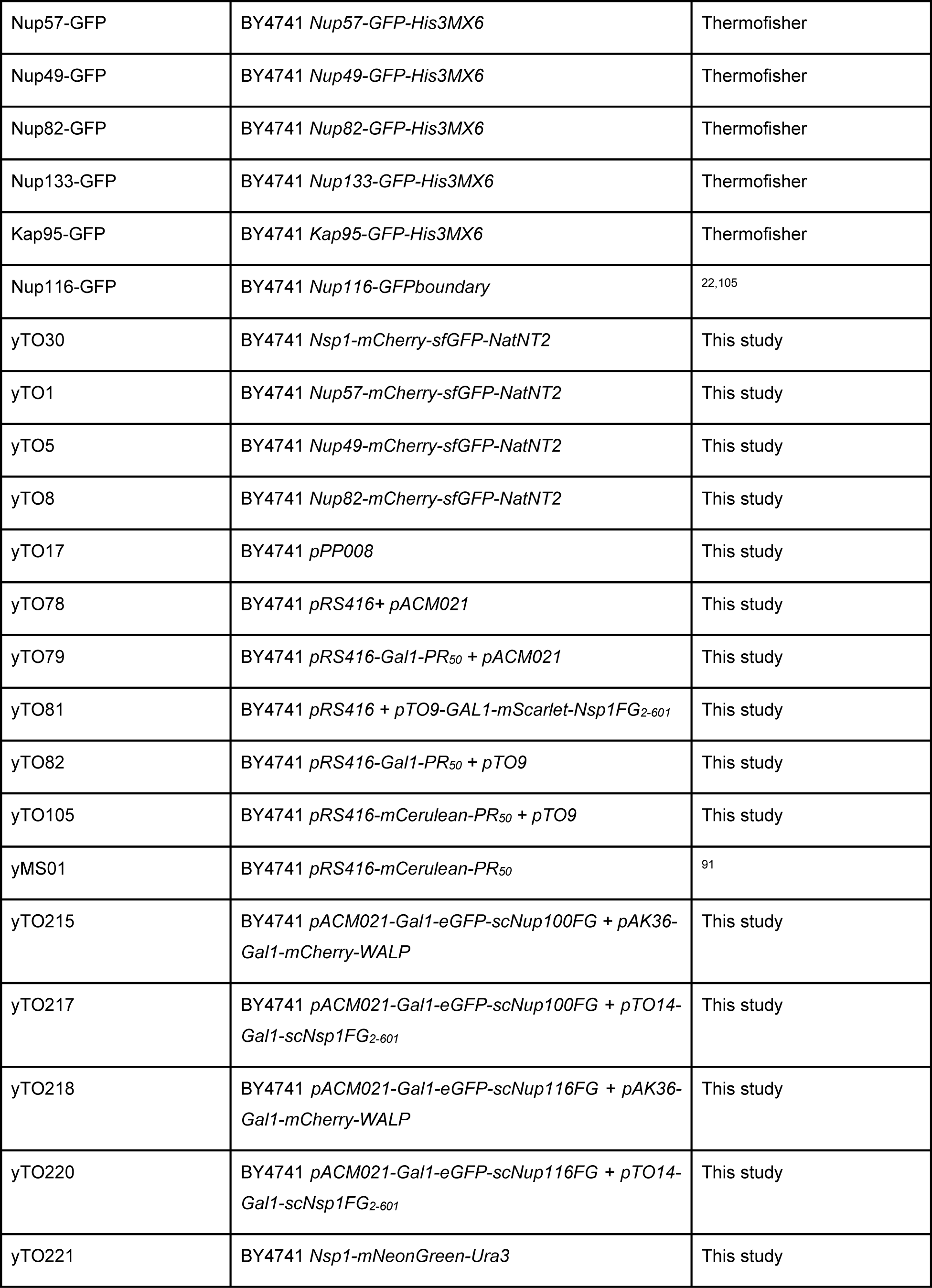
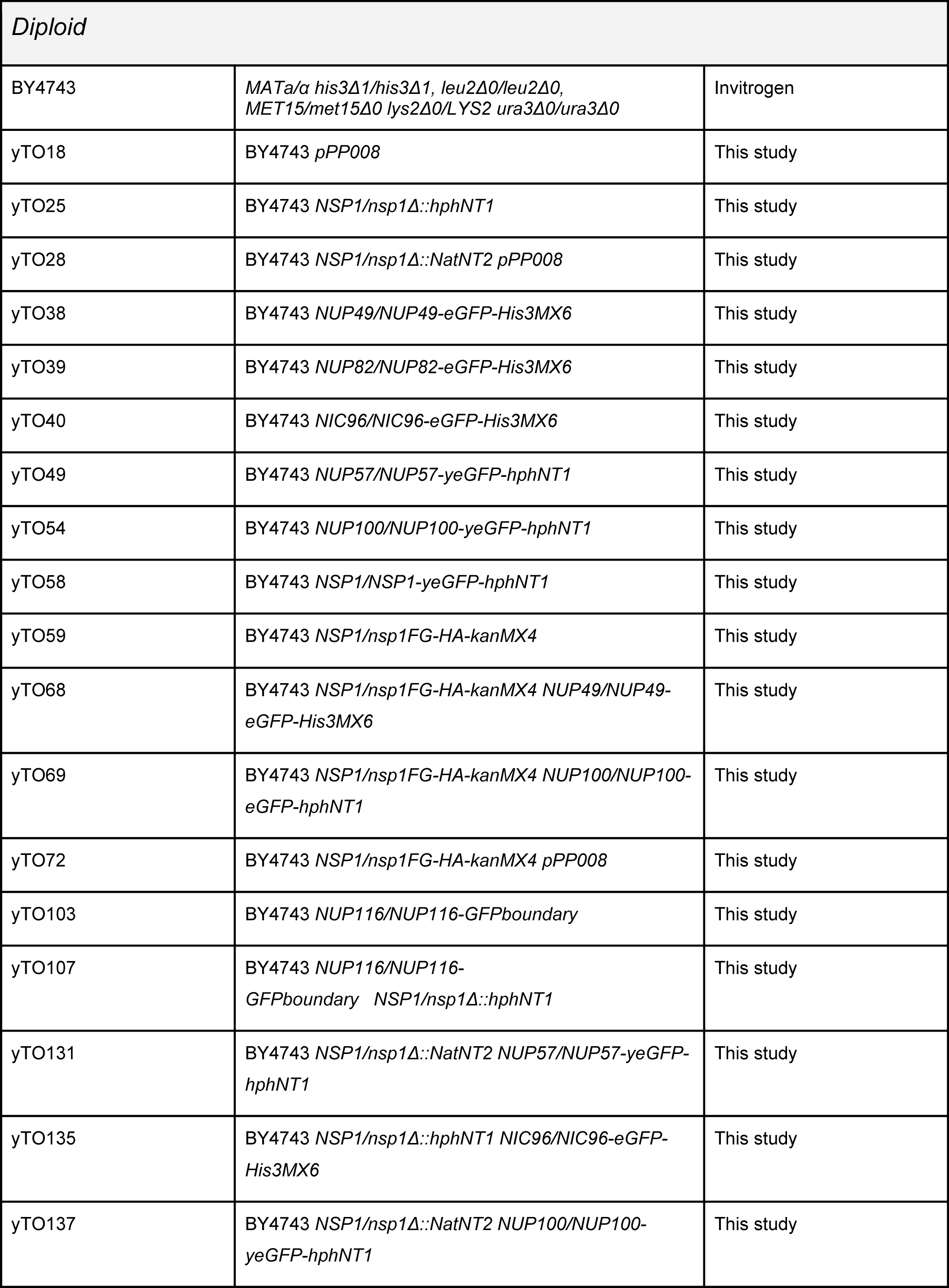
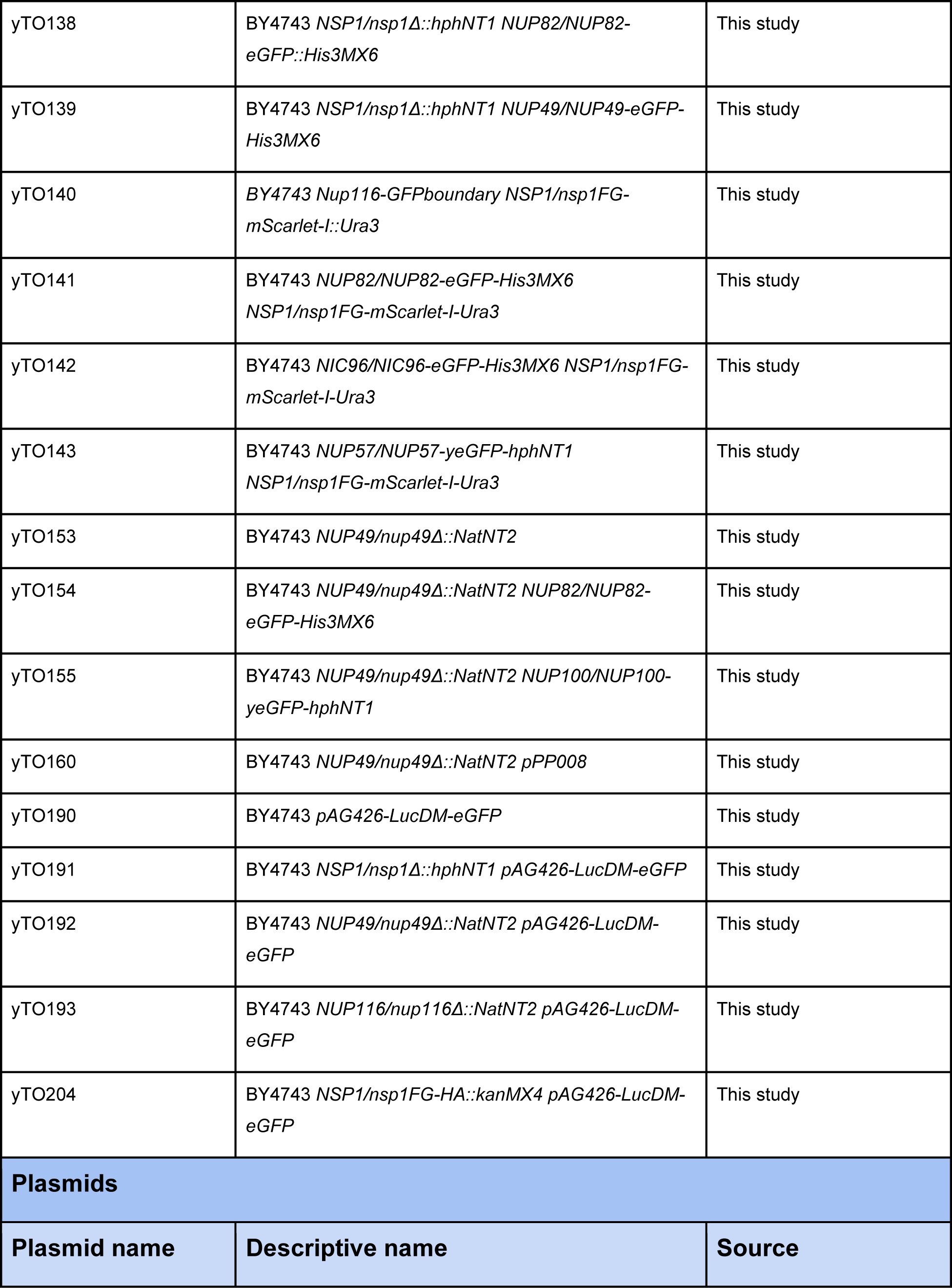
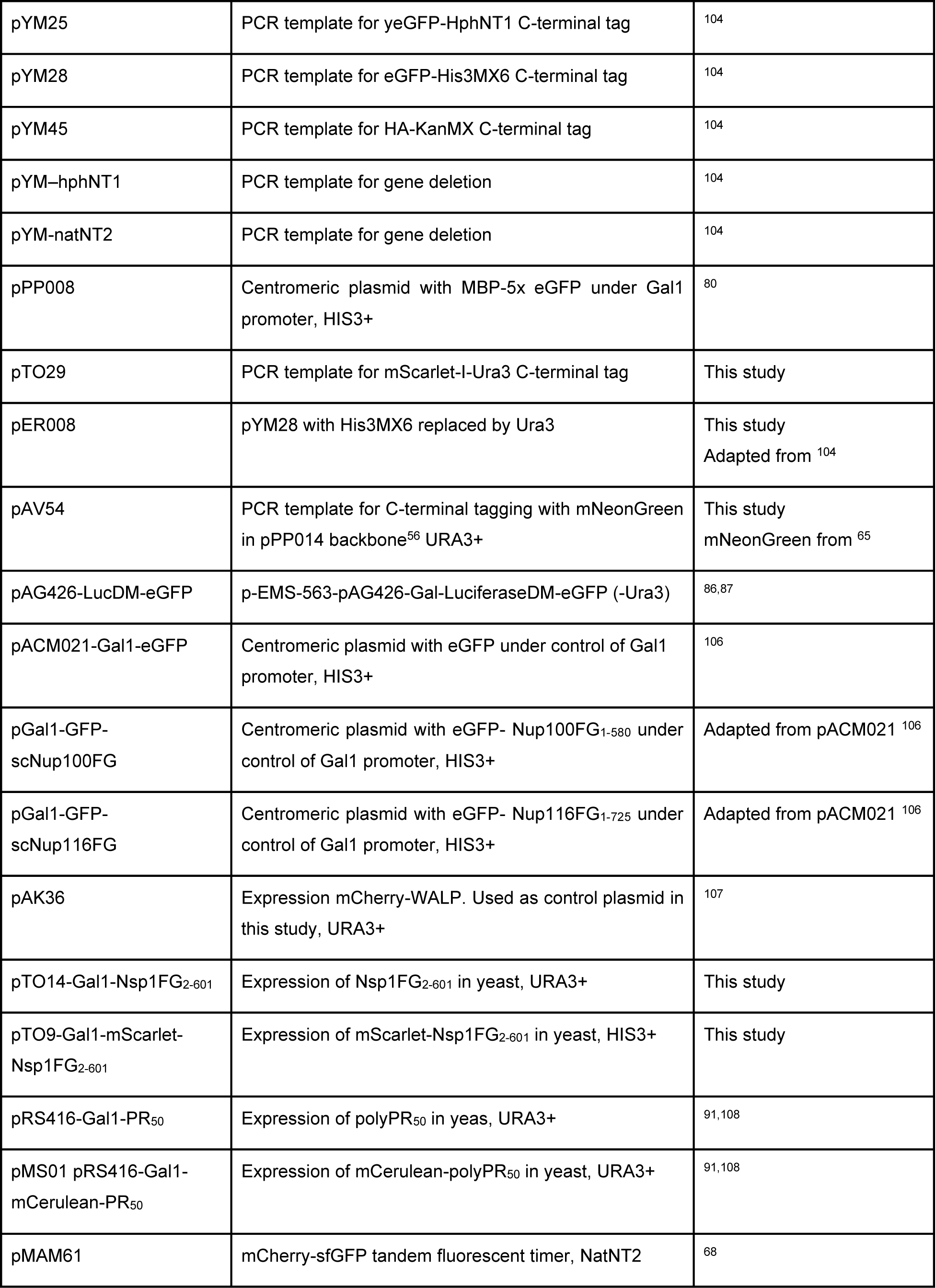
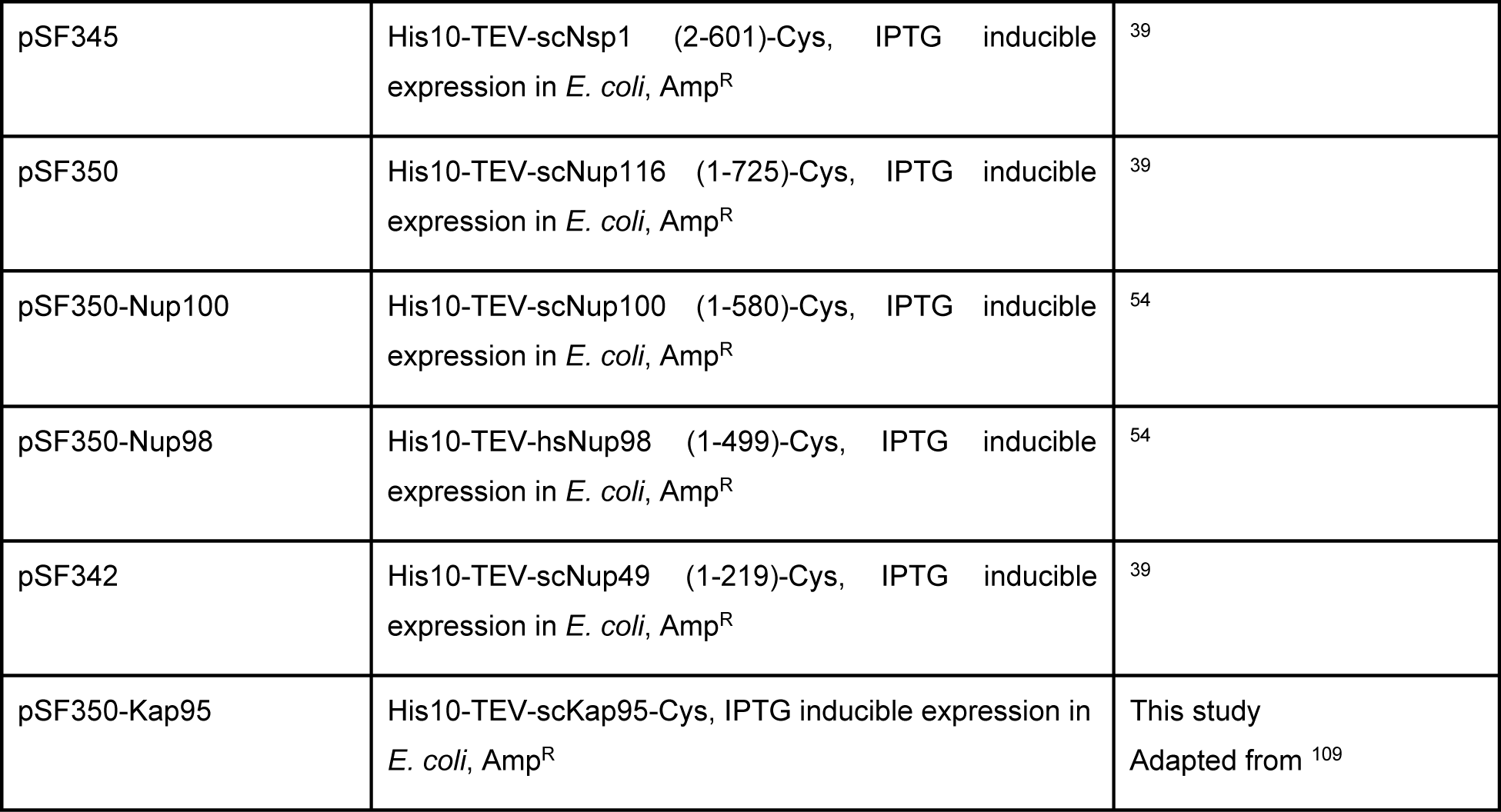

#### Microscopy

Images were acquired at 30°C using a Deltavision Elite (Applied Precision) microscope equipped with an Olympus UPlanSApo 60x or 100x (NA1.4) oil immersion objective using InsightSSIT Solid State Illumination. Detection was performed by either an EDGE sCMOS5.5 or CoolSNAP HQ2 camera. Images were acquired in 25-35 Z-slides and deconvolved using softWoRx software (GE Healthcare). Images were analyzed using Fiji (Image J, National Institute of Health).

#### Microfluidic Chip

Microfluidic chips were used as previously described^66^. Every 20 minutes brightfield images were taken to follow cell division and replicative age. For the experiments with cells expressing Nsp1-mNeongreen a 48-hour timelapse was set up and fluorescent images (20 Z-stacks of 0.2 micron) were taken at the start of the experiment (t0) and after 15h, 30h and 45h. For the polyPR_50_ experiments a 24-hour time-lapse was set up and fluorescent images were taken every 20 min (3 Z-stacks of 0.7 micron). A 1:1 mix of cells expressing eGFP and polyPR and cells expressing mScalet-Nsp1FG and polyPR, or a 1:1 mix of wildtype cells expressing eGFP and cells expressing mScarlet-Nsp1FG was loaded in the chip. Fluorescence was measured at 488 nm (emission 525/48) for control cells expression eGFP and at 594 nm (emission 625/45) for cells expressing mScarlet-Nsp1FG.

#### Batch aging

Cells (10 ml) were grown as described above to OD_600_∼0.5 in SD medium, collected by centrifugation (2500rcf, 5min, 4°C) and washed with PBS. The cells were incubated with 5mg/mL sulfo-NHS-LC-biotin for 30 min while shaking at 800 rpm at room temperature. Excessive biotin was washed away, and cells were recovered for 2 hours in SD medium at 30°C, before incubating with 200 µL Streptavidin-coated magnetic beads for 30 min at room temperature. Streptavidin-coated cells were inoculated in 2L SD medium and grown for 20h, 30 °C, 200 rpm. The cells were subsequently collected by centrifugation (2500rcf, 12 min, 4°C) and resuspended in fresh SD medium. Aged cells were then collected with a magnet. Samples of both young and old cells were stained with Fluorescent brighter (Calcofluor white) to stain budscars, indicative for replicative cell age, before imaging.

#### Spot assay

Yeast cultures were grown overnight in SD medium with 2% raffinose and diluted to OD_600_∼0.5 the next day and grown for one more cell division (OD_600_∼1). Four serial 1:10 dilutions were spotted on plates containing SD with 2% galactose and incubated for 48h at 30 °C. Plates were imaged using a ScanMaker 9800XL scanner (MicroTek International, Inc.).

#### Immunofluorescence

Exponential growing cells (OD∼0.5-0.8) were fixed with 4% paraformaldehyde for 30 min, while rotating at room temperature. The cells were resuspended in a spheroplasting buffer (0.1mM KPO_4_, 500uM MgCl_2_, 1.2M Sorbitol) and spheroplasted with 0.6 mg/mL zymolyase (T20) for 5 min at 37 °C. Spheroplasts were collected by centrifugation (20rcf, 10min, RT) and plated on 50mm Glass Bottom dishes (No. 1.5 uncoated, gamma irradiated, MatTek Corporation) coated with 0.1mg/mL ConcanavalinA for 10 min. For extra permeabilization and flattening of the spheroplast they were incubated with ice-cold methanol for 6 min, ice-cold acetone for 30 sec and quickly dried on a 95 °C heat block. Spheroplasts were blocked with 2% BSA, 0.1% Tween20 in PBS for 1h at room temperature. Primary antibody mouse monoclonal (32D6) anti-Nsp1 was diluted 1:1000 in blocking solution and incubated overnight at 4 °C. Spheroplasts were washed with PBS + 0.1% Tween20. Secondary antibody anti-mouse-AF594 was diluted 1:500 in blocking solution for 2h in the dark at room temperature. Secondary antibody was washed off with PBS and spheroplasts were imaged in PBS directly after washes.

Images were analyzed using Fiji (Image J, National Institute of Health). To identify the regions of interest (ROI), intensity-based thresholds were set for both channels. A manual threshold was set for the anti-Nsp1 signal to create a mask for the Nsp1 foci, and the automatic Otsu threshold was used for the GFP signal to create a mask for the nup or Kap95 particles. The “AND” operation of the ImageJ ImageCalculator plugin was used to create a new mask defining the area of overlap between the two masks. This mask was used to quantify the degree of colocalization/percentage of overlap between the channels within the Nsp1 foci.

#### Proteomic sample preparation

##### Whole cell lysate preparation

For MassSpec whole cell lysates were made from 20 mL yeast cultures of OD_600_∼0.8-1.2. Cells were collected by centrifugation (1811rcf, 5min, 4 °C), lysed in lysis buffer (50mM HEPES pH7.5, 200mM NaOAc pH7.5, 1mM EDTA, 5mM MgOAc, 5% glycerol, 1% TritonX100, 3mM DTT, 1 mM PSMF and 1 tablet protease inhibitor cocktail) and disrupted with glass beads (disruptor beads 0.5mm) in the FastPrep. Whole cell lysates were collected by serial centrifugation and total protein concentrations were determined using the Pierce BCA protein assay kit.

##### Sample preparation for proteomics analysis

The whole cell lysates in presence of SDS sample buffer were run 6 mm into a precast 4-12% Bis-Tris gels containing 50 µg total protein. The band containing all proteins was visualized with Biosafe Coomassie G-250 stain and excised from gel. Small pieces were washed subsequently with 30% and 50% v/v acetonitrile in 100 mM ammonium bicarbonate (dissolved in milliQ-H_2_O), each incubated at RT for 30 min while mixing (500 rpm) and subsequently with 100% acetonitrile for 5 min, before drying the gel pieces at 37 °C. The proteins were reduced with 10 mM dithiothreitol (in 100 mM ammonium bicarbonate dissolved in milliQ-H_2_O, 30 min, 55 °C) and alkylated with 55 mM iodoacetamide (in 100 mM ammonium bicarbonate dissolved in milliQ-H_2_0, 30 min, in the dark at RT). The gel pieces were washed with 100% acetonitrile for 30 min while mixing (500 rpm) and dried at 37 °C before overnight digestion with trypsin (1:100 g/g) at 37 °C. The peptides were eluted from the gel pieces with 75% v/v acetonitrile plus 5% v/v formic acid (incubation 20 min at RT, mixing 500 rpm). The elution fraction was diluted with 0.1% v/v formic acid for cleanup with a C18-SPE column (SPE C18-Aq 50 mg/1mL, Gracepure). This column was conditioned with 2×1 mL acetonitrile plus 0.1% v/v formic acid,and re-equilibrated with 2×1 mL 0.1% v/v formic acid before application of the samples. The bound peptides were washed with 2×1mL 0.1% v/v formic acid and eluted with 2×0.4 mL 50% v/v acetonitrile plus 0.1% v/v formic acid. The eluted fractions were dried under vacuum and resuspended in 0.1% v/v formic acid to a final concentration of around 1 µg/µL total protein starting material.

##### Discovery-based proteomics analyses

Discovery mass spectrometric analyses were performed on an orbitrap mass spectrometer with a nano-electrospray source (Orbitrap Q Exactive 480, Thermo Scientific). Chromatographic separation of the peptides was performed by liquid chromatography on a nano-HPLC system (Ultimate 3000, Thermo Scientific) using a nano-LC column (Acclaim PepMapC100 C18, 75 µm x 50 cm, 2 µm, 100 Å, Thermo Scientific). In general, an equivalent of 1 µg total protein starting material was injected using the µL-pickup method with 0.1% v/v formic acid as a transport liquid from a cooled autosampler (5 °C) and loaded onto a trap column (µPrecolumn cartridge, Acclaim PepMap100 C18, 5 µm, 100 Å, 300 µmx5 mm, Thermo Scientific). Peptides were separated on the nano-LC column using a linear gradient from 2-30% v/v acetonitrile plus 0.1% v/v formic acid in 75 min at a flow rate of 300 nL/min. The mass spectrometer was operated in positive ion mode and data-independent acquisition mode (DIA) using isolation windows of 9 m/z with a precursor mass range of 400-1200, switching the FAIMS between CV −45V and −60V with three scheduled MS1 scans during each screening of the precursor mass range. LC-MS raw data were processed with Spectronaut (version 15.7.220308) (Biognosys) using the standard settings of the directDIA workflow except that quantification was performed on MS1, with a yeast SwissProt database (www.uniprot.org, 6721 entries). For the quantification, local normalization was applied, and the Q-value filtering was set to the classic setting without imputing. For downstream processing Log2 fold changes compared to the wildtype were calculated and plotted and compared with the aging proteomics data set^21^.

##### DotBlot

Whole cell lysates were prepared as described above (section: Whole cell lysate preparation). For the dot blot, 1 µg of total protein diluted in SDS sample buffer was spotted in triplicate on a nitrocellulose membrane (Amersham, Protran 0.2um) in a 96-well biodot microfiltration apparatus (Biorad), following the manufacturer’s instructions. The dot blot membrane with bound proteins was blocked for 4 hrs in TBS-T (50 mM Tris-HCl 150 mM naCl, 0.1% Tween-20, pH 7.4) with 3% BSA. The membrane was then incubated with cMyc antibody in a 1:5000 dilution in 3% BSA in TBS-T for 16 hrs. The membrane was washed three times with TBS-T, and subsequently incubated with HRP-conjugated mouse IgG kappa binding protein (1:5000 in 3% BSA in TBS-T) for 1 hr. The membrane was washed 3 times with TBS-T, twice with TBS and bound antibodies were detected by chemiluminescence using ECL prime (Amersham) in a Biorad chemidoc.

#### Protein purification and LLPS and aggregation assays

Protein purifications of the FG-domains were performed as described in^54^. For Kap95 purification the same protocol was followed with a different buffer (20 mM HEPES/NaOH pH7.5, 110 mM KOAc, 2 mM MgCl2, 0.1% (w/v) Tween20, 10 µM CaCl2 and 1 mM 1,4-Dithiothreitol (DTT)). For expression of Nsp1FG domain in E*. coli* the intron in the starting codon was absent^39^. To assess the effect of Nsp1FG-domain on other FG-domains, purified proteins were diluted in the assay buffer (50 mM Tris–HCl and 150 mM NaCl, pH 8.0) to a protein concentration of 3 µM, unless otherwise indicated. The FG-Nups were incubated in the assay buffer with different concentrations of Nsp1FG-domain (0, 1.5, 3, 6 µM) for 1h at room temperature. All experiments were performed in low protein binding tubes.

For the sedimentation assay, proteins were separated into a soluble and insoluble fraction by centrifugation (17000 rcf, 10 min, RT). Insoluble and soluble fractions were separated with SDS-PAGE and stained with Brilliant Blue G overnight. Band intensities were determined with Fiji (Image J, National Institute of Health) and insoluble fractions were calculated compared to the total protein (insoluble + soluble). For the sedimentation assays only unlabeled purified proteins were used. Filter trap assay was performed as previously described in Kuiper *et al.* (2022)^54^. For the filter trap assay the FG-Nups were used at 3 μM (1:1 molar ratio) and Kap95 was used at either a concentration of 1.5 μM (1:0.5 molar ratio), or 3 μM (1:1 molar ratio).

For microscopy, FG-domains were labeled with fluorescein-5-maleimide as described in Kuiper et al. (2022)^54^. Imaging was performed in a temperature-controlled environment at 25 °C. Fluorescein-5-maleimide-labeled Nup116FG and Nup49FG were mixed with unlabelled Nup116FG and Nup49FG to a final labeling concentration of 5% and 25% respectively. 3 μM of Nup116FG and Nup49FG fragments were diluted out in a droplet of assay buffer on a chambered coverslip (Ibidi, #80826) at 25 °C for 1 or 2 hours, in the absence or presence of unlabelled Nsp1FG, Nup2FG or Kap95. Nsp1FG and Nup2FG were used at a concentration of 3 μM (1:1 molar ratio) and added either from the start (assembling particles) or after 1h (preformed particles). Kap95 was used at a concentration 3 μM (1:1 molar ratio) and added from the start. Within one droplet there was large variability in the properties of Nup116FG particles in the presence of Nsp1FG depending on where in the droplet images were taken. For example, showing surface wetting in the center of the droplet. Therefore, we imaged in 8 different positions across the droplet, to have a complete representation of the particle population within the droplet. Particle properties were analyzed using an ImageJ plugin as described in (Bergsma *et al*, in preparation).

#### Coarse-grained molecular dynamics simulations

Simulations were performed with the coarse-grained 1BPA-1.1 model for IDPs^77–79, 110, 111^. This residue-scale implicit solvent model accounts for the sequence-specific backbone stiffness and discriminates between the 20 amino acids through residue-specific hydrophobic, electrostatic, and cation-pi interactions. The reader is referred to Dekker *et al.* (2022)^79^ for a detailed description of the 1BPA-1.1 model. All simulations are conducted with the GROMACS^112^ molecular dynamics software (version 2019.4) using a Langevin dynamics integrator with inverse friction coefficient ***γ***^-1^ = 50 ps and a timestep of 0.02 ps. The simulation temperature is set to 300 Kelvin, the salt concentration is modeled as 150 mM by setting the Debye screening constant ***κ*** to 1.27 nm^-1^.

The setup of the simulations can be broken up into two parts: the phase separation of the FG-Nup Nup116FG_1-725_ and the addition of Nsp1FG_1-601_ monomers to the phase separated FG-Nup condensates. The FG-Nup phase separation simulation is set up in a similar manner as described in Dekker *et al.* (2022)^54^. To reduce the equilibration time, a pre-build condensate structure is used as the starting configuration for the simulation. This initial condensate structure is obtained by systematically packing 95 Nup116FG molecules in a cubic periodic box at a density of 2 residues per nm^3^. This closed-packed configuration is simulated in NVT for 500 ns to obtain a homogeneous mixture. Then, the box dimensions are increased to obtain the desired concentration of 51 µM for Nup116FG. This system is simulated for 2.5 µs, which was previously found to be sufficient to reach the equilibrium state^79^.

The final coordinates of the Nup116FG equilibration simulation are used as the starting configuration for the addition of Nsp1FG molecules. First, the FG-Nup condensate is centered in the simulation box, after which 140 Nsp1FG monomers are randomly placed around the condensate. Nsp1FG monomer conformations are sampled from a single chain simulation. We made sure there is no overlap between any of the beads by enforcing a minimal distance of 0.5 nm between all chains. The simulation box dimensions are kept constant, resulting in an Nsp1FG concentration of 75 µM (Figure S6A). The Nup116FG + Nsp1FG system (consisting of ∼153000 beads) is simulated over 6 µs (see Figure S6B).

The number of Nup116FG and Nsp1FG molecules in the condensate is determined using the built-in GROMACS utility *gmx clustsize*, where a molecule is considered as being part of the condensate when at least one residue is within 0.7 nm of any residue of the condensate^110^. The intermolecular contacts between Nup116FG and Nsp1FG in the condensate are analyzed every 1 ns over the final 2 µs of the trajectory. For each trajectory frame, the specific molecules that are part of the condensate are identified (using the *gmx clustsize)*, for which a contact matrix is computed using MDAnalysis^113, 114^. Also here, two residues are considered to be in contact when their separation is less than 0.7 nm. The average number of intermolecular contacts is then obtained by summing all individual contact matrices and normalizing this with the respective number of Nup116FG and Nsp1FG in the condensate. Finally, the contact map is time-averaged over the trajectory.

#### Statistical analysis

Biological and technical replicates are indicated in the figure legends. Data was tested for normality by the Shapiro-Wilk test. For normally distributed data an unpaired t-test or ordinary one-way ANOVA with Šídák’s multiple comparisons test was used. Otherwise, a nonparametric Mann Whitney test or Kruskall-Wallis test with Dunn’s multiple comparisons test was performed to indicate statistical differences. For fraction of cells with foci a Chi-square test was performed. Graphs and statistical analyses were generated using Prism9 software (GraphPad).

